# Systematic review of residual toxicity studies of pesticides to bees and comparison to language on pesticide labels using data from studies and the Environmental Protection Agency

**DOI:** 10.1101/2023.06.05.543089

**Authors:** Leah K. Swanson, Andony Melathopoulos, Matthew T. Bucy

## Abstract

**BACKGROUND:** Residues of pesticides on crops can result in mortality to foraging bees. The likelihood of mortality can be mitigated by applying pesticides in the evening so that their residues dissipate by the following morning when bees resume foraging. The dissipation rates of different pesticides, or their residual toxicity, is captured in a public-facing database compiled by the U.S. Environmental Protection Agency (EPA), but the database includes only a fraction pesticides bees are likely to encounter in the environment. Pesticide applicators in the U.S. encounter a Pollinating Insect Hazard Statement on pesticide labels, which coarsely indicate which products dissipate over the course of an evening. There is reason to suspect that these statements may not align with residual toxicity data, given previous findings of significant misalignment from published data discovered on the acute toxicity section of the Pollinating Insect Hazard Statement. Without a complete database of residual toxicity estimates, however, it is not possible to determine whether the residual toxicity components of the Pollinating Insect Hazard Statement similarly diverge from published studies.

**RESULTS:** We compiled 48 studies and calculated the residual time to 25% mortality (RT_25_) of each assay for three different bee species (*Apis mellifera, Nomia melanderi*, and *Megachile rotundata*). Our findings were compared to the EPA published database of RT_25_ values. Of the RT_25_ values that we could compare, we found that over 90% of the values support a similar conclusion to EPA: that the active ingredient has extended residual toxicity (i.e., residues cause greater than 25% mortality for eight hours or more). Next, we compared our values and the EPA’s values to the Pollinating Insect Hazard Statement in the Environmental Hazards sections of 155 EPA registered product labels. Of these labels, a little less than a third (27%) presented their residual toxicity in a manner inconsistent with their calculated RT_25_ and current EPA labeling guidelines. Moreover, over a third (33%) of labels contained an active ingredient which was neither listed under EPA’s RT_25_ database nor had a published study to estimate this value.

**CONCLUSION:** Residual toxicity of pesticides is a key parameter used by pesticide applicators to reduce impacts of their applications to bees. We provide the first evidence that many pesticide labels may convey residual toxicity information to applicators that is not correct and could lead to bees being exposed to toxic residues on plants. We also show large gaps in the availability of contemporary residual toxicity study for many pesticides, suggesting either researchers should conduct studies to estimate RT_25_ values for these products, or EPA should make data from registrants more readily available. Finally, our analysis identified significant variation found between RT_25_ values among different bee species tested, and different formulations of the same active ingredient, suggesting these factors should be incorporated into future bee residual toxicity studies.

## Introduction

Pesticides can have negative impacts on individual bees and bee colonies when toxic products are applied to blooming plants that are bee attractive (Botías *et al*., 2017; Chauzat *et al*., 2010; Kiljanek *et al*., 2017; Tosi *et al*., 2018; Graham *et al*., 2021). Bees can become exposed to pesticide residues when foraging on pesticide-treated plants, which can result in their mortality if the residues are at levels that are acutely toxic to them. Mortality, however, may be lessened if the pesticide is applied in the evening, when bees are not foraging. This allows for an interval over which the pesticide can dissipate on the plant sufficiently to become relatively non-toxic to bees when they resume foraging the next day. Evening pesticide applications as a way to mitigate exposure, however, is predicated on the assumption that the residues of the pesticides will weather sufficiently before bees resume foraging the next morning and come into contact with treated leaves and flowers (Barmaz *et al*., 2010; Fischer and Moriarty, 2011; Honey Bee Health Coalition, 2019; Smodiš Škerl *et al*., 2009).

The rate at which acute toxicity of pesticides to bees dissipate from plant surfaces is known as the pesticide’s residual time. Pesticide registrants in the U.S. are required to estimate the residual time for all pesticides that contain one or more active ingredients that is acutely toxic to bees (*i.e.,* acute contact toxicity lethal dose to 50% of the honey bees (LD_50_ is less than 11 micrograms of pesticide per bee) and the use pattern indicates that bees are likely to be exposed (40 CFR 158.630(d)). EPA provides guidance for registrants on how to conduct a trial to estimate residual time (United States Environmental Protection Agency [USEPA], 2012a). These trials involve spraying a field crop (typically alfalfa) with a pesticide, allowing residues to weather for set intervals, then harvesting plant material and placing it in a cage with honey bees (*Apis mellifera*). The bees are free to walk over the plant material for a set period of time (typically 24 h), after which the number of dead bees is counted. Residual time is expressed as the weathering interval after which the mortality of bees contacting the foliage falls reaches 25% mortality (referred to as the residual time to 25% mortality or RT_25_). The basic pattern of these trials pre-dates EPA guidelines and have been used by toxicologists since the 1960s (*e.g.*, Wiese, 1962).

A key threshold residual time identified by EPA is known as extended residual toxicity. A pesticide with extended residual toxicity is one that cannot be applied safely in the evening as residues would cause more than 25% mortality of bees in a cage assay. Although the residual time threshold is not specified in EPA’s Label Review Manual (2012), elsewhere EPA indicates that a pesticide with extended residual toxicity has RT_25_ > 8h (Office of Chemical Safety and Pollution Prevention [OCSPP], 2012). Typically, pesticide labels in the U.S. only indicate whether pesticides that are acutely toxic to bees have extended residual toxicity or not and generally do not list RT_25_ values (OCSPP, 2012).

RT_25_ is an important tool in determining how to best mitigate the risk of bee exposure to pesticides residues. The importance of RT_25_ estimates for pesticide applicators when selecting and applying a pesticide is evinced by state Cooperative Extension publications that list RT_25_ values from published studies (*e.g*., Hooven *et al*., 2016). Furthermore, RT_25_ estimates are used by EPA in order to characterize the hazards and risks of pesticides to pollinating insects. The EPA requires that a product’s residual toxicity to bees be communicated on the product label in a way that is reflective of the RT_25_ value. EPA has produced guidance for their reviewers and pesticide registrants on the language they will typically suggest for different RT_25_ values (USEPA, 2012b). This information will typically be available in the Environmental Hazards section of the label, but it is not federally enforceable and is used as an informational tool for pesticide applicators (USEPA, 2012b). However, pesticide labels rarely state the RT_25_ value, so this information is not readily accessible to pesticide applicators, crop advisors or extension educators, there remains a demand for better guidance on the dissipation rates of bee toxic products under field conditions.

Notably, more recent EPA guidance (EPA, 2017) would provide more specific mitigation language around the extended residual toxicity threshold for the safe application of pesticides during bee pollination. These new guidelines provide federally enforceable specific use instructions for residual toxicity stating that if extended residual toxicity (residues persisting for greater than 8 hours *i.e*., extended residual toxicity) is not present for a pesticide it can be applied 2 hours before sunset when pollinators are least active (EPA, 2017). Pesticide registrants have begun adopting this guidance, one example is Harvanta 50SL (Summit Agro^TM^, Durham, NC, EPA registration number 71512-26-88783), which states for fruiting vegetables (Crop Group 8-10) “foliar application of this product is prohibited to a crop from onset of flowering until flowering is complete unless the application is being made in the time period between 2 hours prior to sunset until sunrise.” While this shows that some labels have been written in accordance with this new policy, many pesticide labels still follow pre-2017 guidance in communicating residual toxicity to bees (e.g., Product Dursban 50W, EPA Registration Number 62719-72; Product Merit 2F, EPA Registration Number 432-1312). In addition to the 2017 guidance, EPA released a public summary of RT_25_ estimates compiled from registrant-submitted data to the public (EPA, 2014). Notably, the summary only included studies that have “undergone quality assurance reviews to ensure that the data are scientifically sound”, and, in turn, is missing several widely used active ingredients (*e.g*., bifenthrin). Regardless, the omissions pose a challenge to researchers looking to compare pesticide label language on residual toxicity to RT_25_ values.

There is a need for investigating pesticide label language against studies that characterize environmental risks. Bucy and Melathopoulos (2019), for example, found that roughly 32% of pesticide labels analyzed had at least one error in the communication of acute toxicity to bees, or the adverse effects caused after a short exposure time to an active ingredient (OCSPP 850.3000). These authors, however, were unable to do a similar analysis with residual toxicity statements because of the absence of a comprehensive database of RT_25_ values.

Our objective was to provide the first analysis of pesticide label statements communicating residual toxicity to bees in comparison to actual RT_25_ values. We approached the challenges experienced by Bucy and Melathopoulos (2019) by creating a database of RT_25_ values to compare to pesticide label statements. Our approach to creating a database was to assemble all published residual toxicity studies and characterize variability in methodologies used to assess residual toxicity. We then conducted a meta-analysis to calculate RT_25_ estimates for each pesticide and validated these estimates against values published by EPA (EPA, 2014). We used the validated database to analyze the residual toxicity statement on pesticide labels and to determine how RT_25_ estimates vary by the rate of pesticide used, the formulation of the pesticide, and bee species.

## Materials & Methods

### 2.1 Selection of studies

We located putative residual toxicity studies using Web of Science with the search term “residual toxicity” as well as the names of bee taxa commonly used in residual toxicity assays: “*Apis*”, “*Nomia*”, “*Megachile*”, “*Bombus*” and bumble bees. This search returned a total of 130 studies. Next, we located residual toxicity studies on the alfalfa leafcutting bee (*Megachile rotundata*) from proceedings of the Western Alfalfa Seed Growers Association (2004-2017), resulting in an additional 17 studies. We also evaluated series of Bee Research Investigation and Integrated Pest and Pollinator Investigation reports released by Washington State University amount to 28 reports. Finally, we obtained 8 studies directly from Bayer CropSciences. In sum, we evaluated 183 residual toxicity studies. We narrowed these studies down to 50 papers by excluding papers that met any of the following criteria: (1) the study was not a primary source of data (*e.g*., review paper); (2) bees were exposed to the pesticide applied to paper instead of on plant material; and/or, (3) the study focused on residual toxicity of pesticides applied to plots but did not involve harvesting plant tissue for caged bees. These criteria were designed to ensure that we only included studies whose residual toxicity methodology broadly followed those of EPA (2016). We removed two additional papers because the author indicated that it was likely that some live bees in the assay were mistakenly counted as dead (Johansen *et al*., 1981) and because the actual active ingredient of the product used was not specified (Walsh, 2011).

### 2.2 Evaluation of studies

This analysis consisted of residual toxicity studies where a pesticide was foliar applied onto a specific crop, and the plant material (*e.g*., foliage, flowers) was harvested at varying time intervals after application. The plant material was then collected and placed in cages with adult bees to contact for 24 hours or longer. Residual toxicity was calculated from variation in bee mortality for bees exposed to plant material harvested at different intervals of weathering. Although studies were selected based on their broad adherence with the methodology developed by EPA (2016), they varied across several test parameters. We categorized the variance of EPA methodology across four key study parameters (Table 1).

**Table 1.**
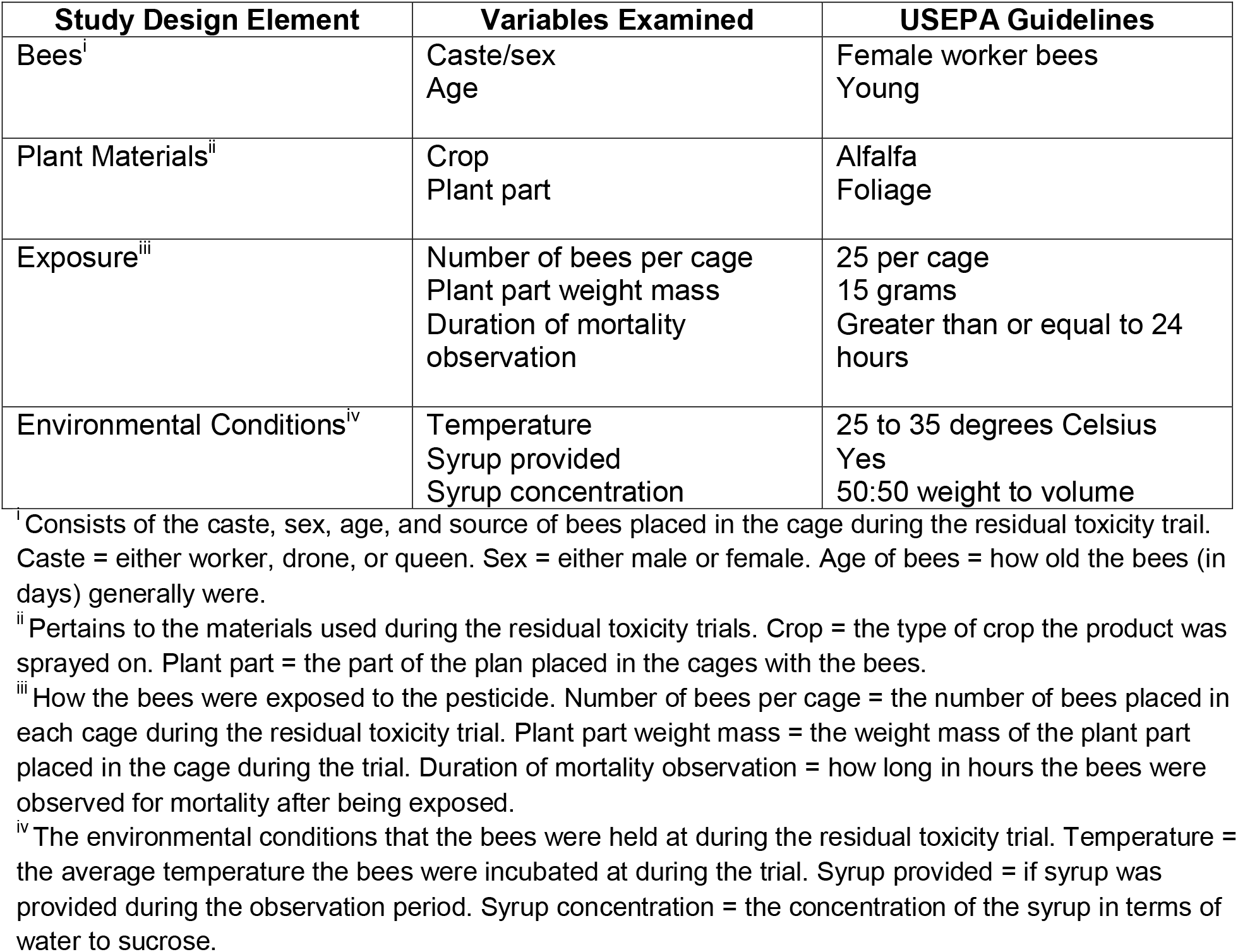
Descriptions of Study Design Elements Examined during the Meta-Analysis

| Study Design Element | Variables Examined | USEPA Guidelines |
| --- | --- | --- |
| Bees<sup>i</sup> | Caste/sex<br>Age | Female worker bees<br>Young |
| Plant Materials<sup>ii</sup> | Crop<br>Plant part | Alfalfa<br>Foliage |
| Exposure<sup>iii</sup> | Number of bees per cage<br>Plant part weight mass<br>Duration of mortality<br>observation | 25 per cage<br>15 grams<br>Greater than or equal to 24<br>hours |
| Environmental Conditions<sup>iv</sup> | Temperature<br>Syrup provided<br>Syrup concentration | 25 to 35 degrees Celsius<br>Yes<br>50:50 weight to volume |

<sup>i</sup> Consists of the caste, sex, age, and source of bees placed in the cage during the residual toxicity trail. Caste = either worker, drone, or queen. Sex = either male or female. Age of bees = how old the bees (in days) generally were.
<sup>ii</sup> Pertains to the materials used during the residual toxicity trials. Crop = the type of crop the product was sprayed on. Plant part = the part of the plan placed in the cages with the bees.
<sup>iii</sup> How the bees were exposed to the pesticide. Number of bees per cage = the number of bees placed in each cage during the residual toxicity trial. Plant part weight mass = the weight mass of the plant part placed in the cage during the trial. Duration of mortality observation = how long in hours the bees were observed for mortality after being exposed.
<sup>iv</sup> The environmental conditions that the bees were held at during the residual toxicity trial. Temperature = the average temperature the bees were incubated at during the trial. Syrup provided = if syrup was provided during the observation period. Syrup concentration = the concentration of the syrup in terms of water to sucrose.

We used the following approaches to standardize methodologies across studies. EPA uses the word “young” to describe the optimal age of bees for residual toxicity trials. We interpreted “young” to mean newly emerged adult (eclosed) bees that were less than 1 day of age (Winston, 1991). Furthermore, when a study reported a range for a parameter, such as for number of bees per cage or temperature, we used the average calculated from the low and high points of the range. In reference to the diet that the bees were fed during the assay, one paper reported the syrup concentration as 91:1 (wt:wt) which we assumed was 1:1 (Mayer, 2001).

We evaluated whether parameters in studies aligned with USEPA recommendations, by counting each testing parameter as described in Section 2.2. We noted whether studies had test parameters that corresponded to those recommended by USEPA or if there was not enough information to determine correspondence.

### 2.3 Calculation of RT_25_ values

Independently of each study’s calculated RT_25_ value, we determined RT_25_ values from each study using percent bee mortality in cages from different time periods. Trials within studies were compiled by active ingredient, the duration residues were allowed to weather, the formulation of the pesticide (emulsifiable concentrate, wettable powder, etc.), application rate, and species of bee used in the cage assay. We removed any trials with only a single weathering period, because these could not be used to calculate residual time. We also removed studies if they did not specify application rates, or percentage mortality and if mixtures of active ingredients were used. If the study provided a percentage mortality for bees in the cage assay, but not the total number of bees used in the cage, we assumed there were a total of 100 bees for the purposes of analysis. USEPA includes both *M. rotundata* and *N. melanderi* as well as *A. mellifera* in their published RT_25_ values but not *Bombus.* Consequently, we also removed *Bombus* trials from this analysis since comparison to USEPA would have been impossible.

We used R Statistical Software (v4.1.1; R Core Team, 2021) with the package Tidyverse (Wickham *et al*., 2019) to calculate the RT_25_ values using regression models where time was the independent variable and percent mortality the dependent variable. We checked for overdispersion in all assays. If the data was not over dispersed, we then calculated the RT_25_ values through a binomial logistic regression. If the data was over dispersed, we calculated the RT_25_ values using a quasibinomial logistic regression.

### 2.4 Comparison of RT_25_ values

We compared RT_25_ values for bee species across active ingredient, formulation, and application rate. *M. rotundata* and *N. melanderi* RT_25_ values were compared to *A. mellifera* RT_25_ values since, currently, EPA generally only requires registrants to conduct residual toxicity assays for *A. mellifera* when applying for product registration. In doing so, we were able to determine whether Environmental Hazards language reflects the RT_25_ estimates of *M. rotundata* and *N. melanderi*.

We validated the database created from calculated RT_25_ values (Section 2.3) by comparing residual times for each active ingredient by application rate, formulation of the product (i.e., emulsifiable concentrate, wettable powder, etc.), and species of bee in the database published by EPA (EPA, 2014). Instead of comparing the RT_25_ estimates themselves, we compared how each database would categorize a pesticide as having extended residual toxicity or not. For example, if our calculated RT_25_ value for a pesticide was less than 6 hours (*i.e*., no extended residual toxicity) and EPA indicated the RT_25_ value was greater than 12 hours (*i.e*., extended residual toxicity), we deemed the two as sufficiently different. Furthermore, we assumed EPA database estimates were accurate. If extended residual toxicity determinations matched those of pesticides from our meta-analysis, this would mean that we could rely on RT_25_ estimates for active ingredients that did not appear in the EPA database. In contrast, if there were substantial misalignment among extended residual toxicity determinations between our calculated and USEPA RT_25_ estimates, we would conclude that our calculation methods significantly differed from EPA’s and our estimates would need to be reevaluated.

### 2.5 Label language analysis

We created a composite database of RT_25_ values from EPA (EPA, 2014) supplemented with calculated values based on the findings of Section 2.4. To determine if RT_25_ values correspond with residual toxicity statements under the Environmental Hazard section of pesticide labels, we used an existing database of residual toxicity statements on pesticides labels developed by Bucy and Melathopoulos (2019) and compared it to RT_25_ values in our composite RT_25_ database. We excluded labels from this analysis if: (1) the Environmental Hazards indicated the product was not ‘toxic’ or not ‘highly toxic’ to bees. This would mean that the active ingredient has an LD_50_ for bees of greater than 11μg/bee, in which case USEPA would not have required that the registrant assess the residual toxicity of that product and/or (2) the product was unlikely to result in exposure to bees (*e.g*., granular formulations). We only used *A. mellifera* RT_25_ values in this analysis since residual toxicity language on pesticide labels is specific to this species (EPA, 2012b). Similar to Bucy and Melathopoulos (2019), We interpreted pesticides without extended residual toxicity RT_25_ times (less than 8 hours) as corresponding to the statement: “Do not apply… while bees…**are actively foraging** the treatment area” and those with extended residual toxicity RT_25_ values (greater than 8 hours) if accompanied with the statement: “Do not apply…if bees…**are foraging** the treatment area” found in EPA label language guidance (USEPA, 2012b). We compared the RT_25_ values to the residual toxicity statements on labels for the same formulation (*e.g*., emulsifiable concentrate, wettable powder, etc.) of the same active ingredients between the calculated RT_25_ values and the pesticide.

The following assumptions were made regarding the interpretation of slight variation from EPA guidance when reviewing labels. “Bees are least active” was interpreted as “Do not apply…while bees…**are actively foraging** the treatment area” and “Bees may forage” was interpreted as “Do not apply…if bees…**are foraging** the treatment area.” If no label language associated with RT_25_ values was present on the label, the data from that label was included as “N/A” in analysis. If no acute toxicity language was present on the label suggesting a LD_50_ of greater than 11 μg/bee, we excluded the active ingredient from our analysis. If an active ingredient had an LD_50_ greater than 11 μg/bee, EPA would not require an acute or residual toxicity statement on the product label.

We determined misalignment between the Environmental Hazards and RT_25_ estimates based on the extended residual toxicity threshold (see Section 2.4). For example, if a label suggested an RT_25_ value less than 8 hours but the database indicated an RT_25_ estimate that was greater than 8 hours, we deemed the label language as not aligning. Many labels had language that corresponded with RT_25_ values but neither EPA nor the literature examined during the meta-analysis had information on the active ingredient in the product or a similar formulation to compare the two. These labels were included in the analysis as labels that had “RT_25_ values missing.”

## Results

### 3.1 Methodology of residual toxicity trials

Almost three quarters of the studies (70%) analyzed used EPA’s recommended leaf foliage as the treated plant material placed in cages during the residual toxicity trials, with around a quarter of the studies using other materials such as flowers. In all studies, bees stocked in cages with treated plant material were fed sucrose syrup *ad libitum*. A majority of studies (69%) aligned with EPA recommendation for a 50% (wt:wt) sucrose to water solution (Figure 1). The temperature at which bees were incubated during the residual toxicity test varied greatly among the studies. Most studies incubated bees outside the temperature range of 25-35°C as recommended by USEPA (Figure 1), tending to incubate at cooler temperatures. The crop used in studies was evenly distributed between EPA recommendation of alfalfa and other crops. The studies that did not use alfalfa used, in descending order of frequency, used cotton, white clover, strawberry, and sunflower. About half of the studies (48%) reported that there were 25 bees placed in the cage for each residual toxicity trial as recommended by EPA, with remaining studies ranging from 10-106 bees per cage. On average, trials using *A. mellifera* had more bees (56) per cage compared to *M. rotundata* (24 bees per cage) and *N. melanderi* (20 bees per cage). The age of the bees used during the residual toxicity trial was mostly uniform across studies with only a quarter of studies using an older age of bees (> 1 day old) than recommended by EPA.

**Figure 1:**
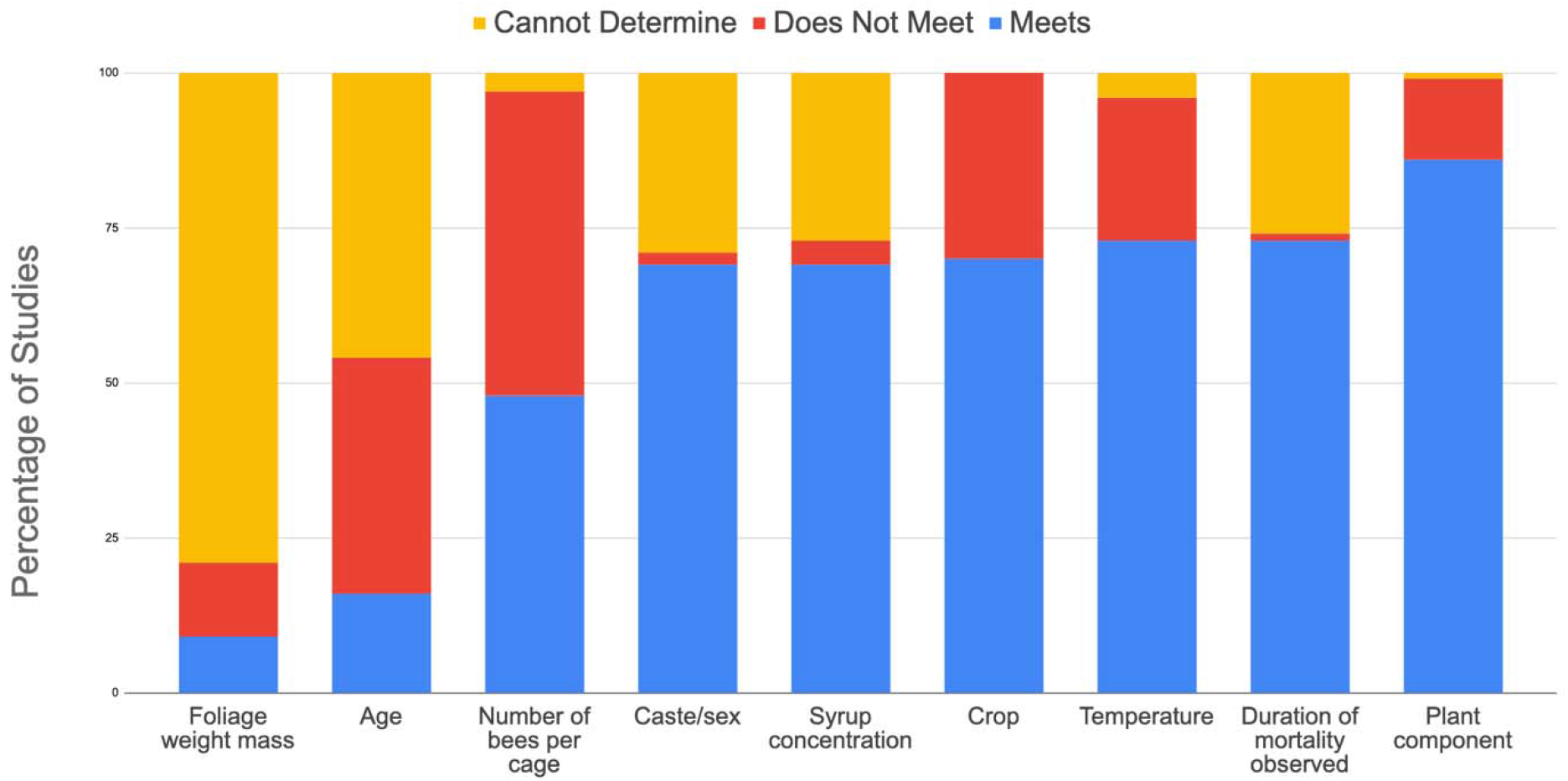
Comparison of methodological parameters of residual toxicity studies (n=48) with percentage of studies that meet USEPA residual toxicity criteria (USEPA, 2012a), do not meet, and cannot determine based on lack of information provided in the study.

### 3.2 RT_25_ calculations and comparisons

We calculated RT_25_ values from 135 of 490 trials in the 48 studies that were reviewed. We were unable to calculate RT_25_ values for the other 355 trial because published mortality percentage values were either above 25% for all time periods reported or were below 25% for the duration of the assay. In these cases, we indicated the RT_25_ value as greater than the longest reported period or less than the shortest reported period respectively.

When comparing RT_25_ values across different formulations, there were six cases where different formulations with the same rate of the same active ingredient resulted in different RT_25_ values (Table 2). These cases were acephate, dimeothate, fipronil, formetanate hydrochloride, naled and trichlorofon. For the same application rates of the same formulation of the same active ingredient, 20 active ingredients had different RT_25_ values with higher application rates having typically longer RT_25_ values. Finally, there were 21 cases where different species (*A. mellifera, M. rotundata,* and *N. melanderi*) resulted in different calculated RT_25_ values even though they were exposed to residues of an active ingredient that was applied as the same formulation, at the same application rate, and allowed to weather for the same amount of time. The most variation in RT_25_ times was seen in active ingredients with the formulation of emulsifiable concentrate with *M. rotundata* consistently having longer RT_25_ times (9 cases) compared to *A. mellifera* (Figure 2).

**Figure 2:**
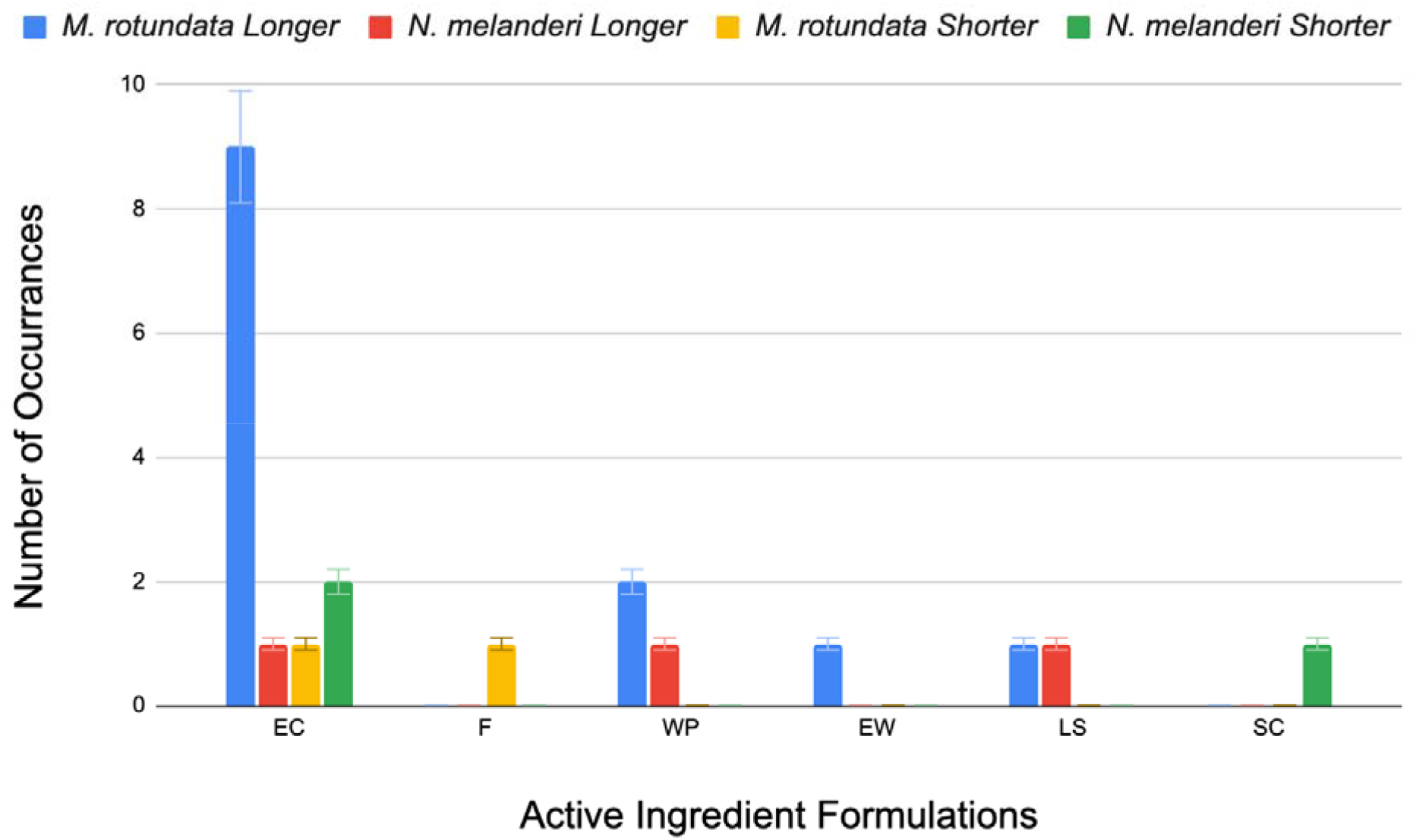
Comparison of bee species across different active ingredient formulations, EC (emulsifiable concentrate), F (flowable), WP (wettable powder), EW (emulsion in water), LS (liquid soluble), and SC (soluble concentrate. *M. rotundata* and *N. melanderi* RT_25_ values were compared to *A. mellifera* and reported as either (1) longer than *A. mellifera* values (“*M. rotundata* longer” and “*N. melanderi* longer”) or (2) shorter than *A. mellifera* values (“*M. rotundata* Shorter” and “*N. melanderi* Shorter”).

**Table 2.**
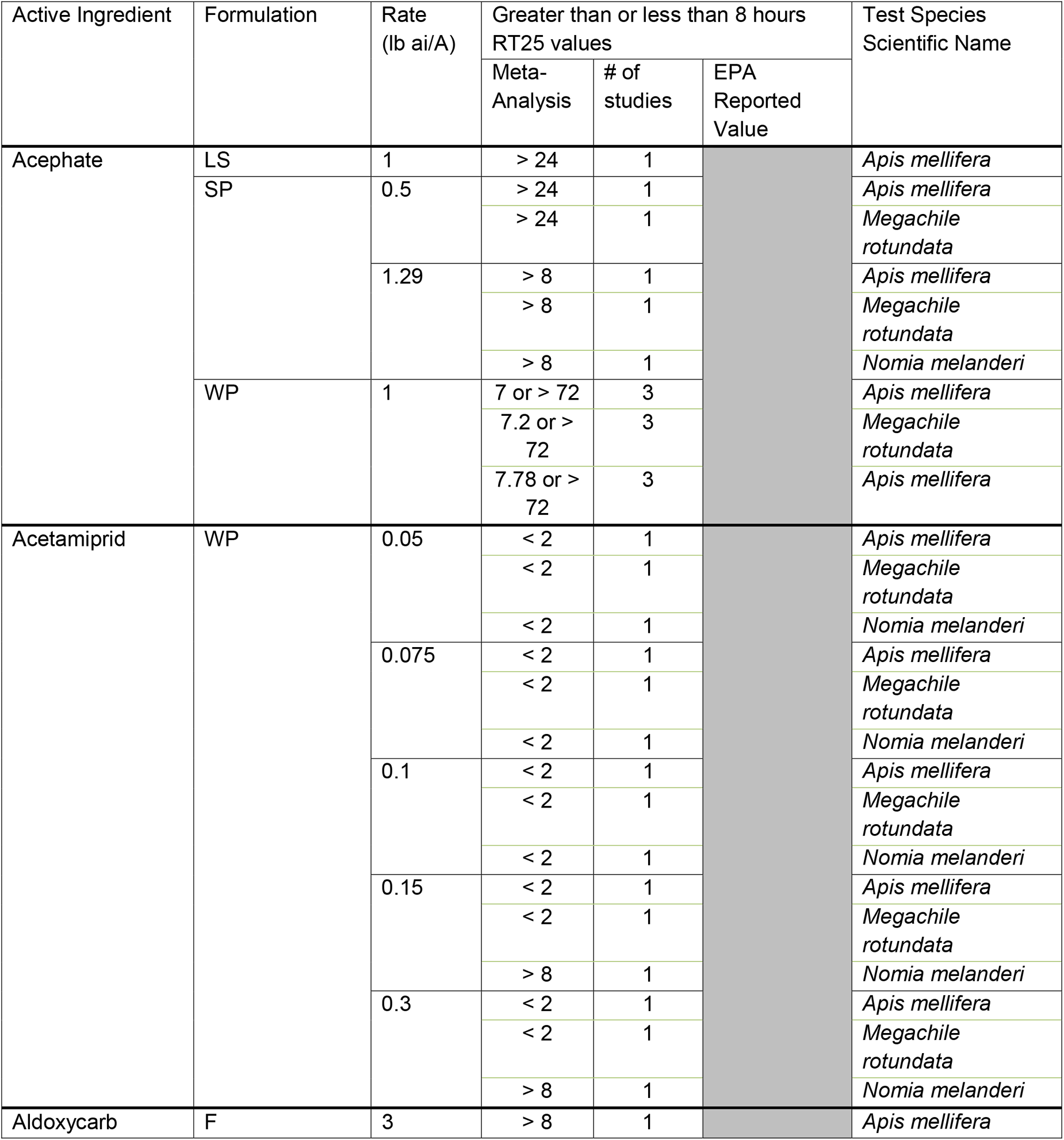
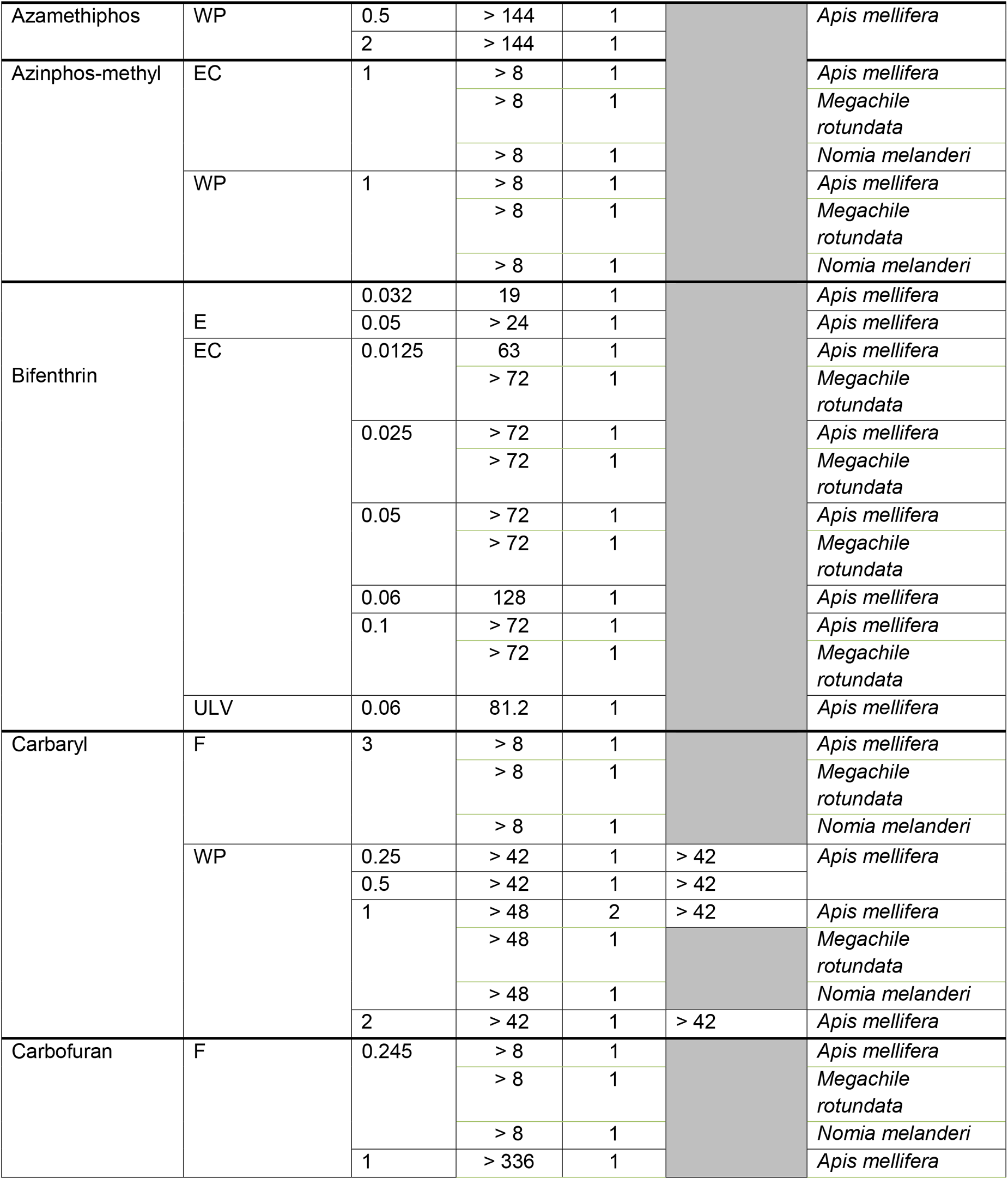
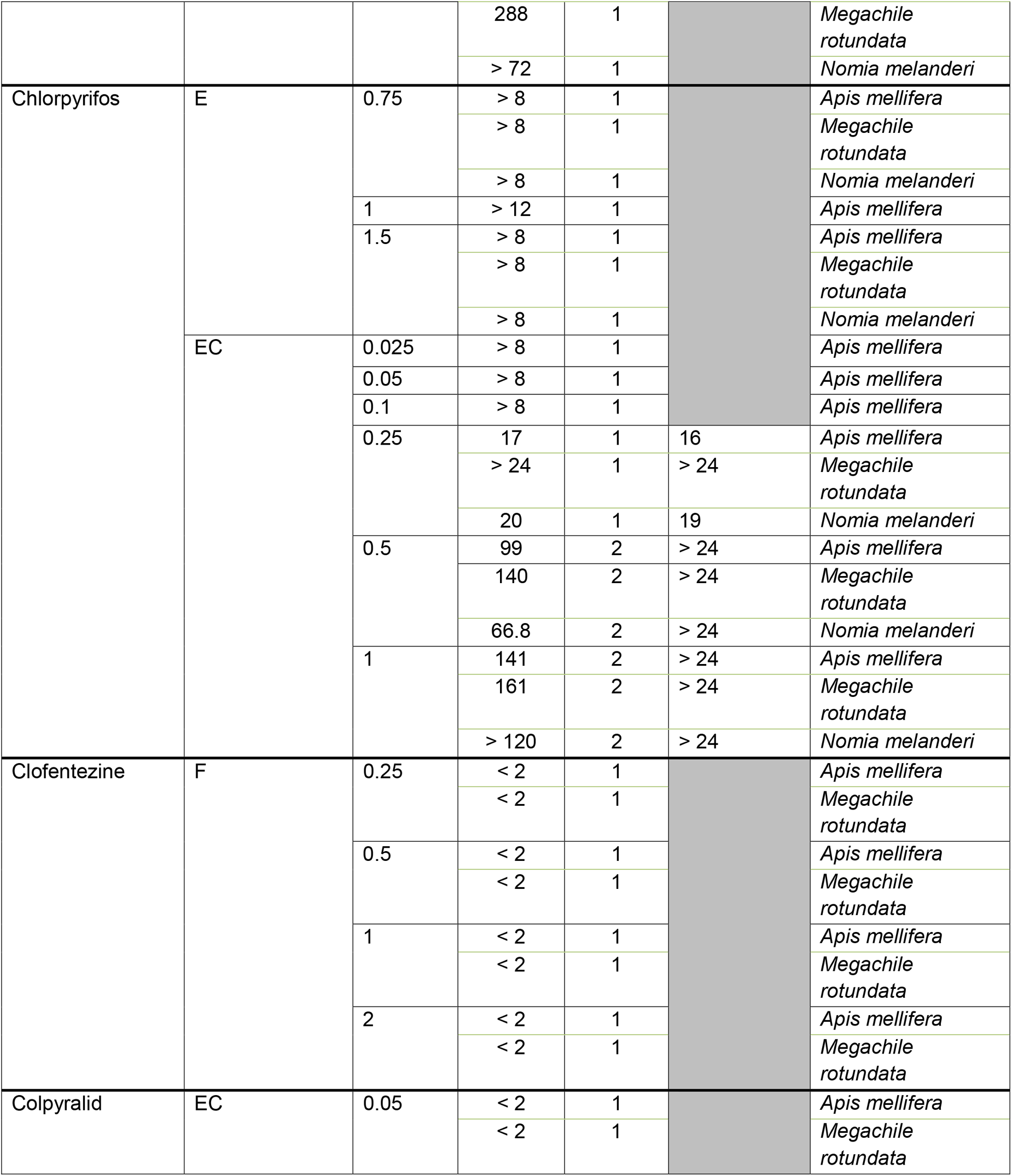
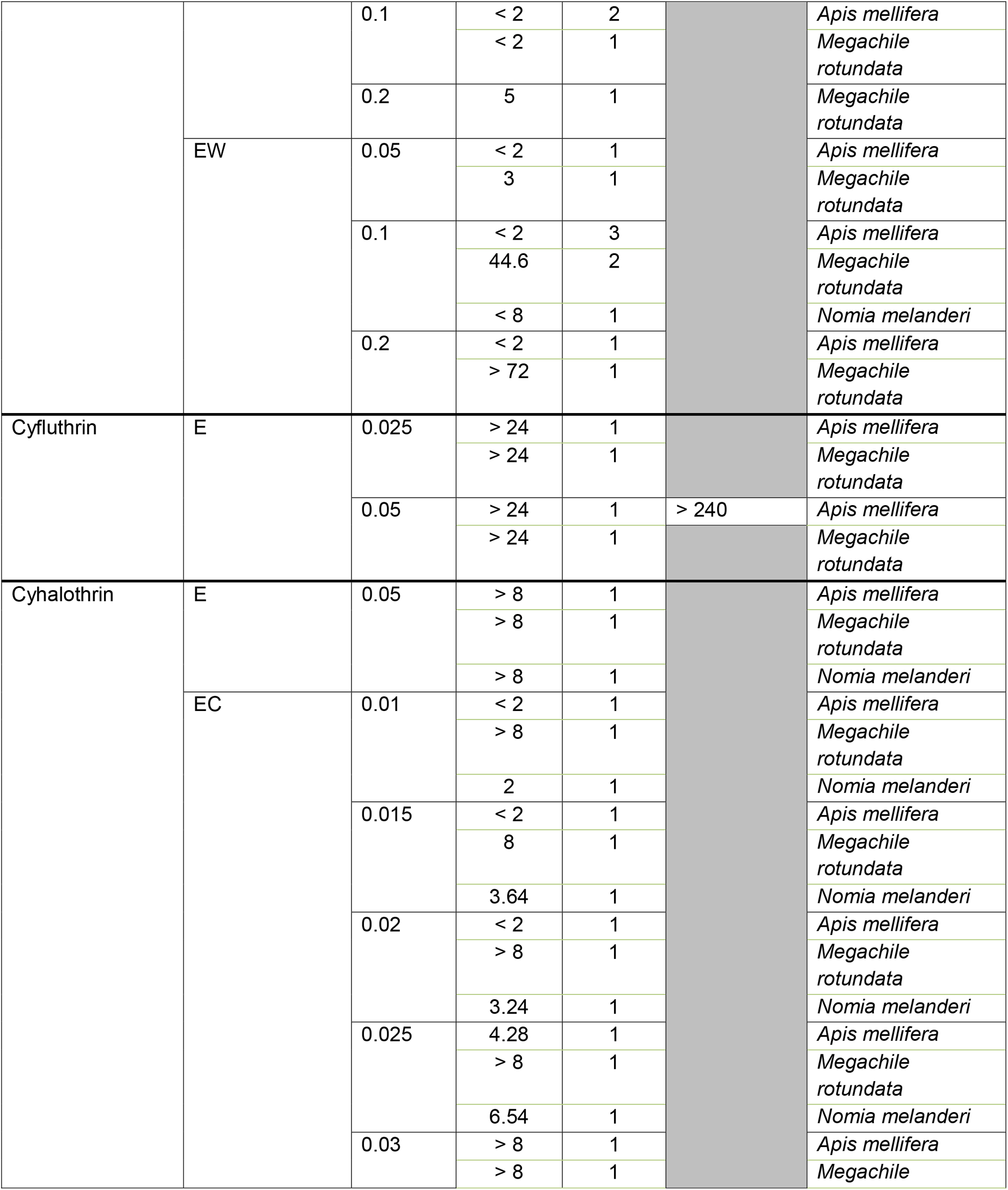
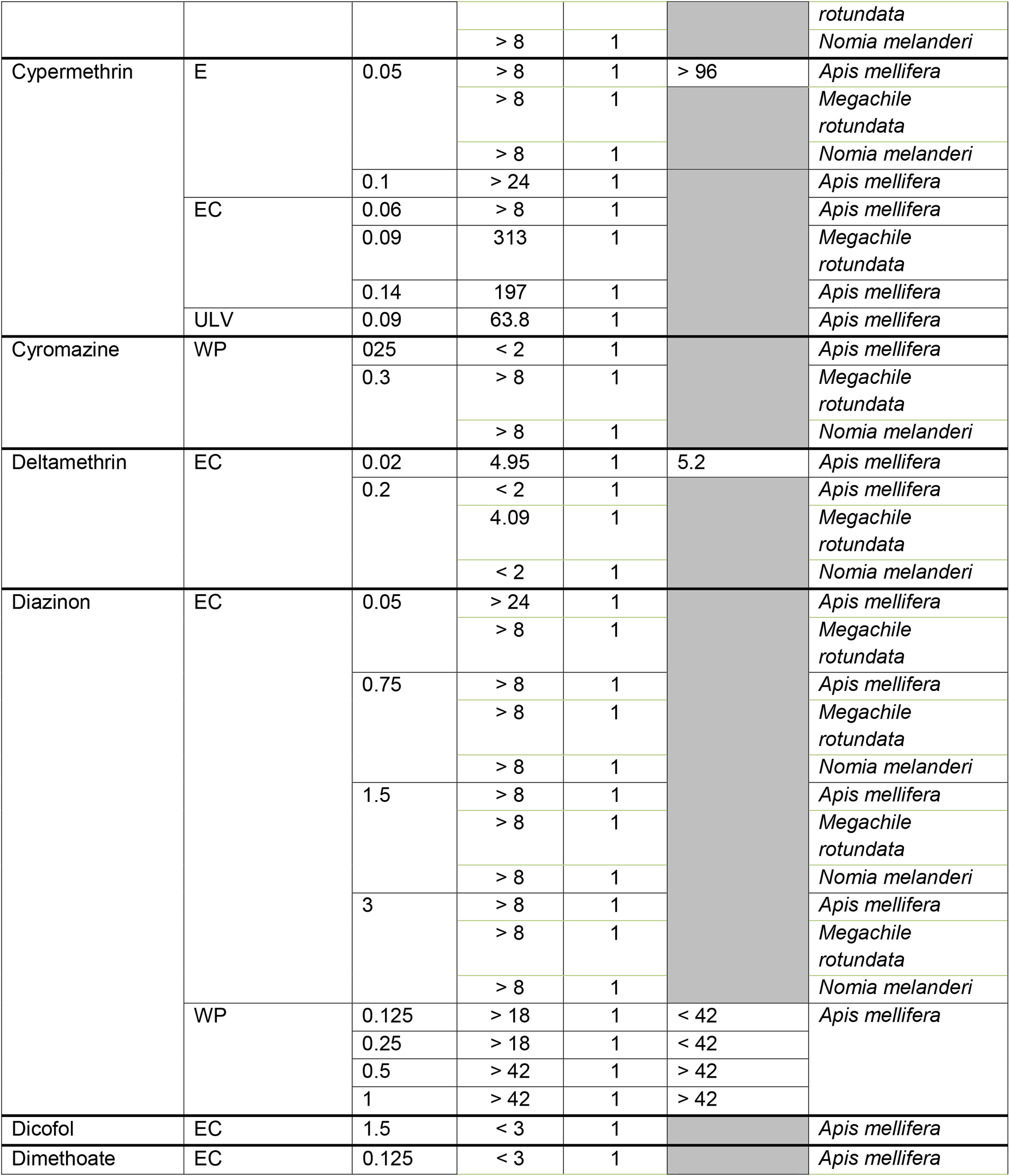
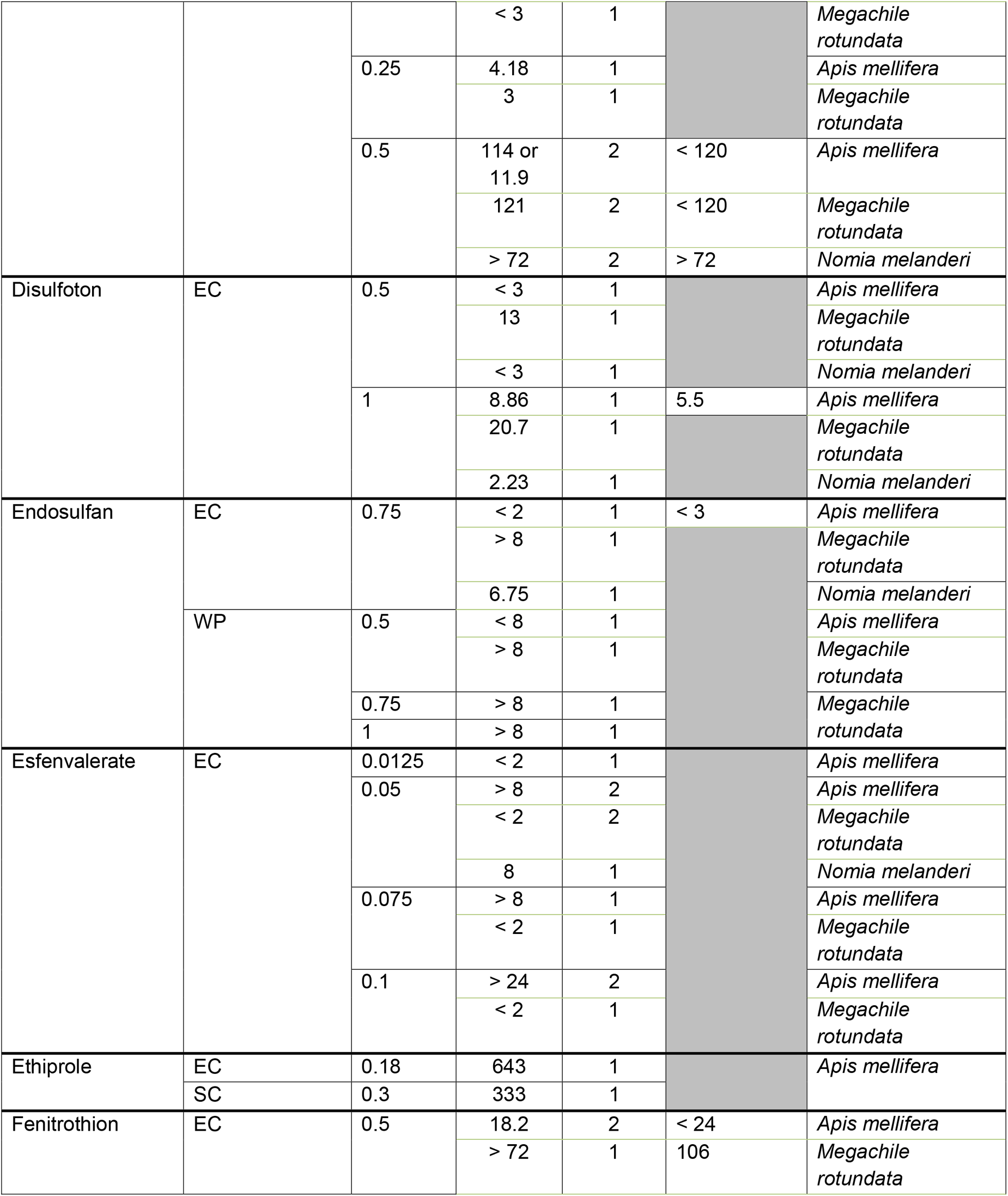
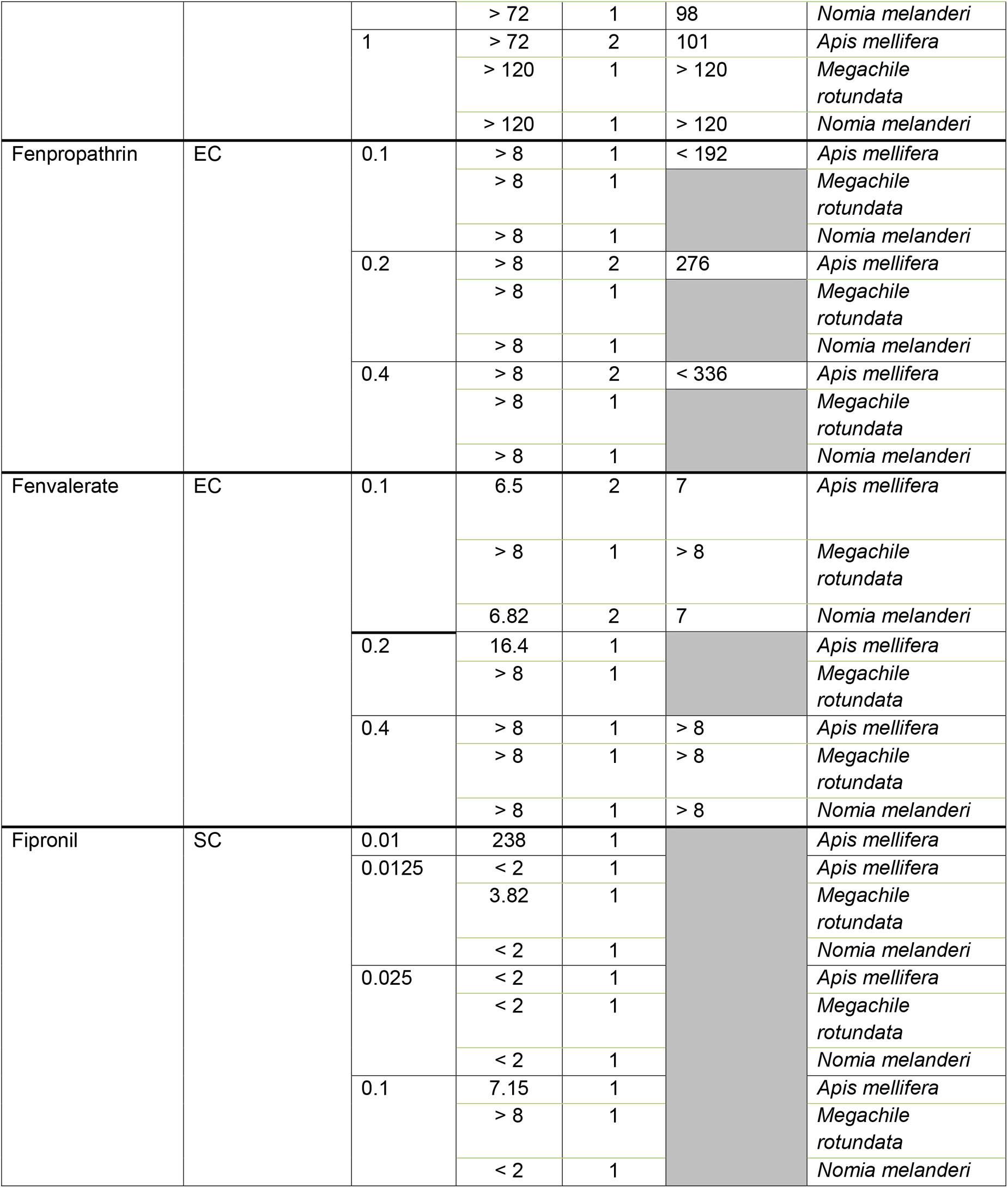
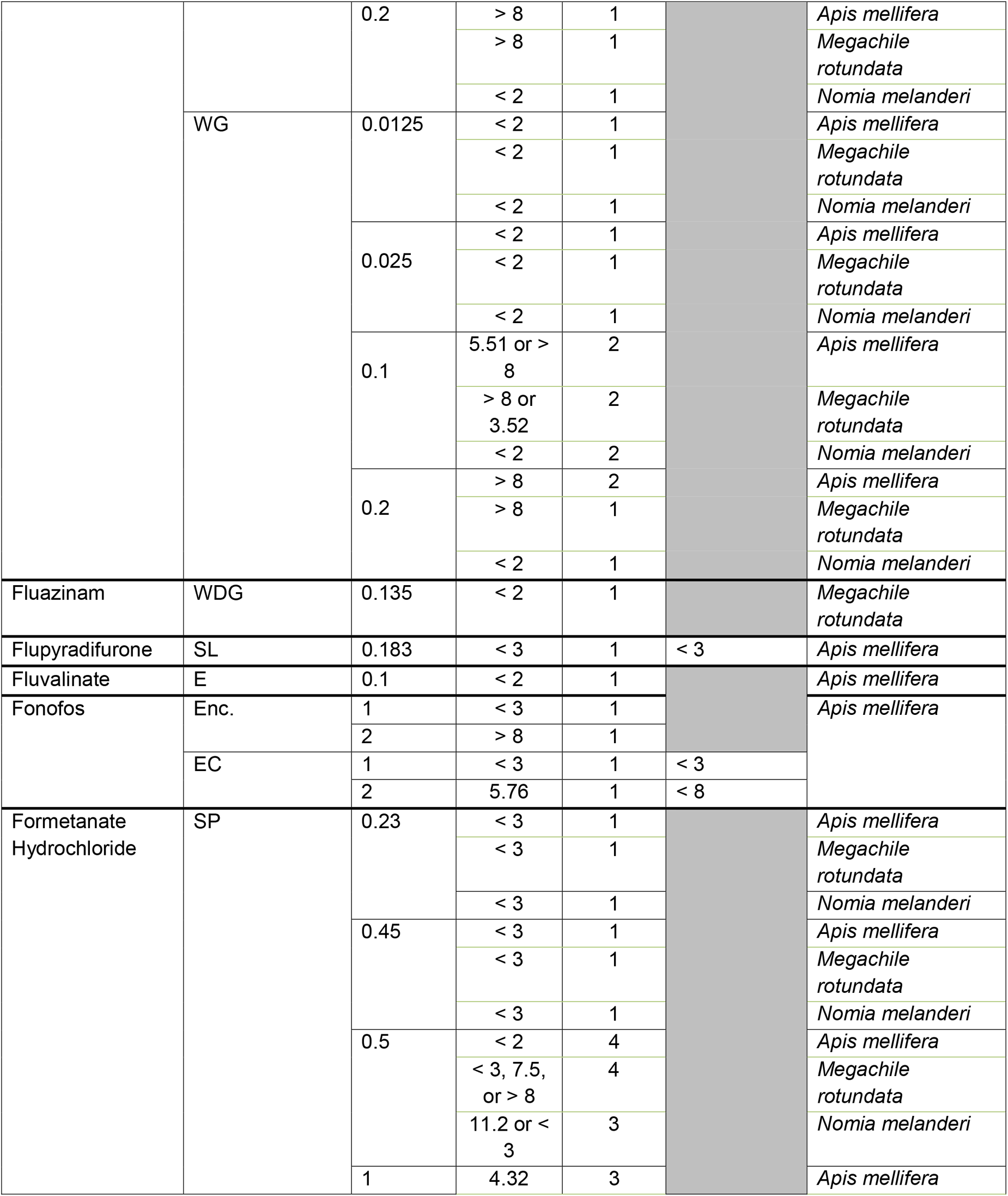
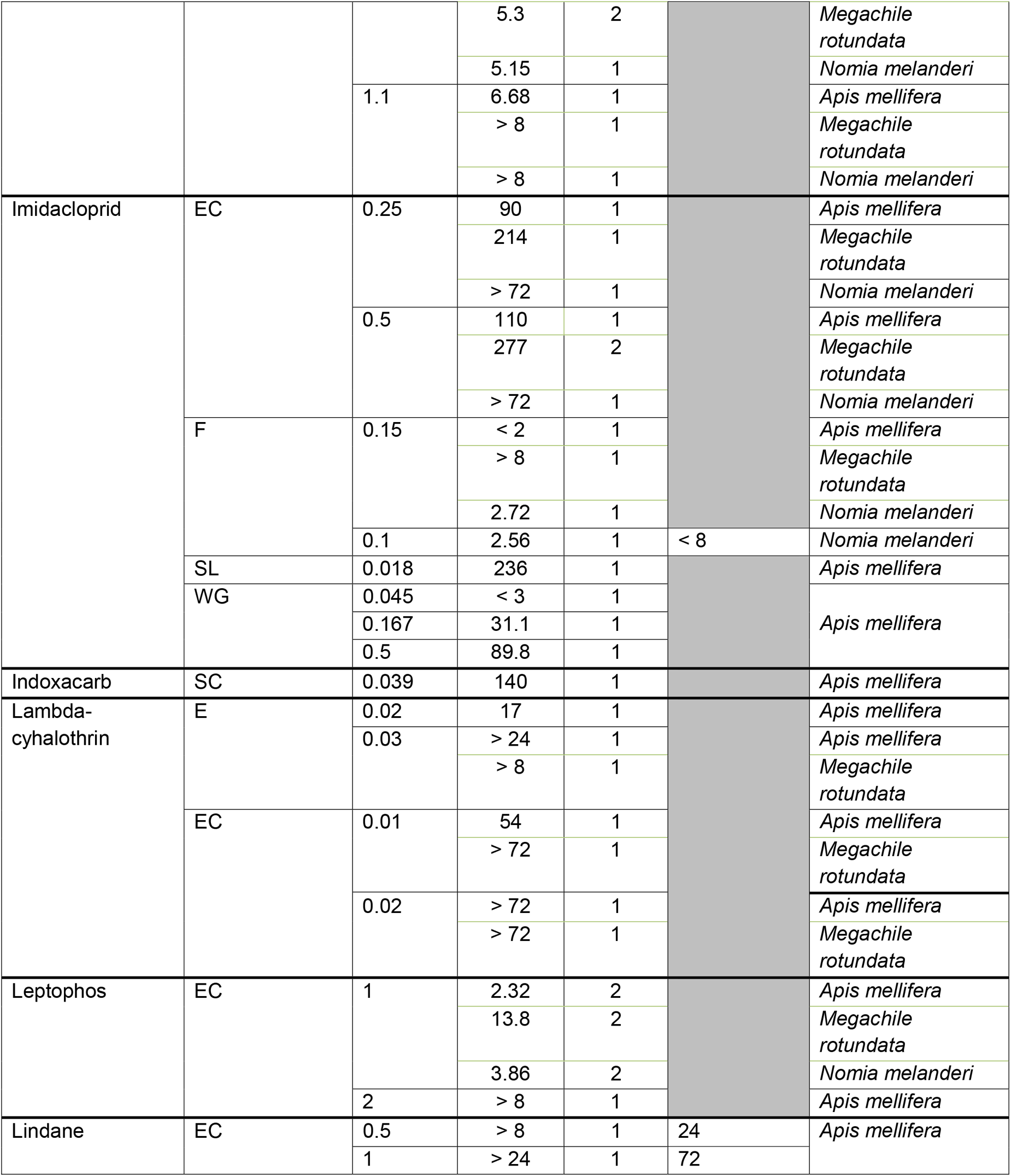
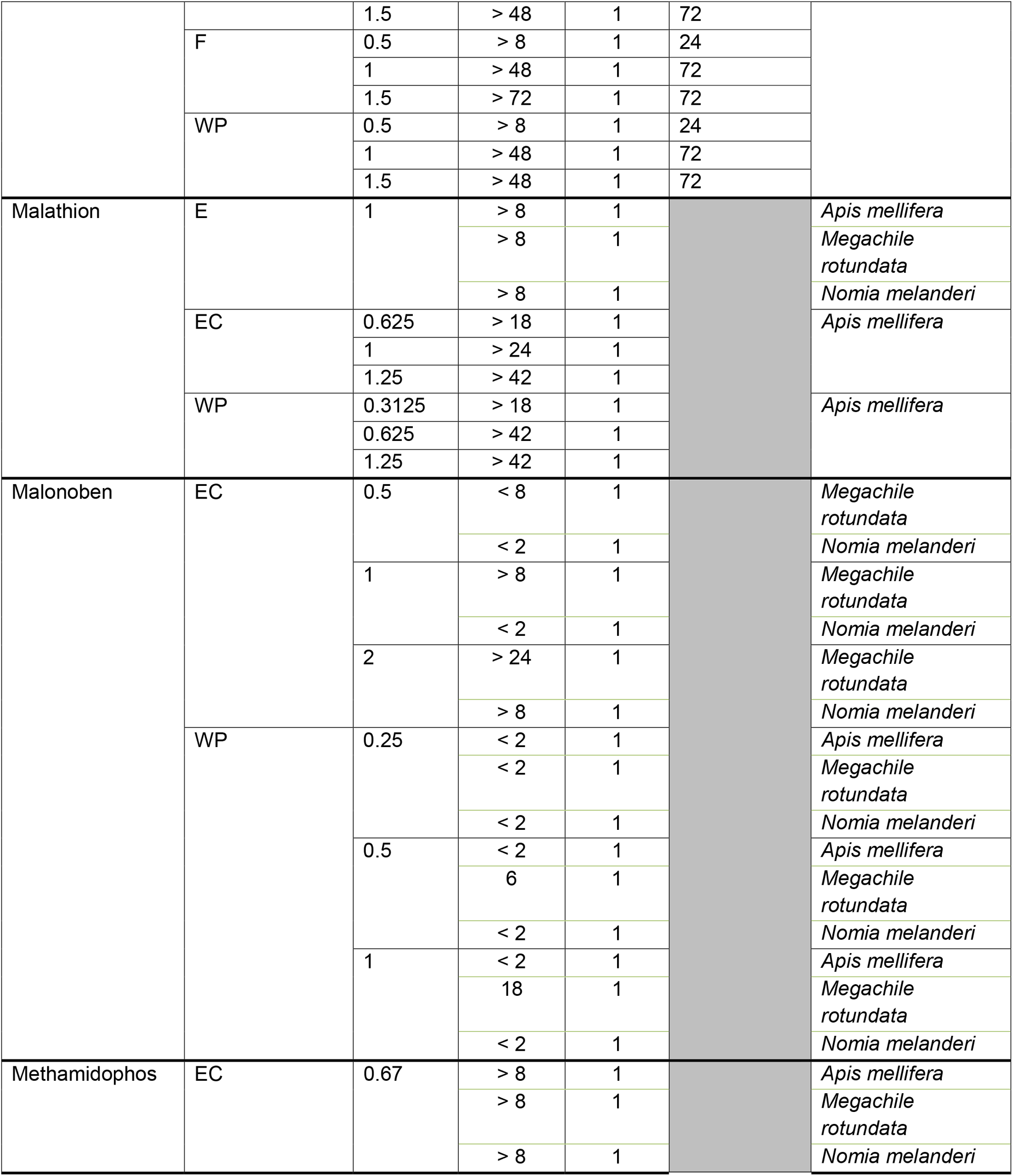
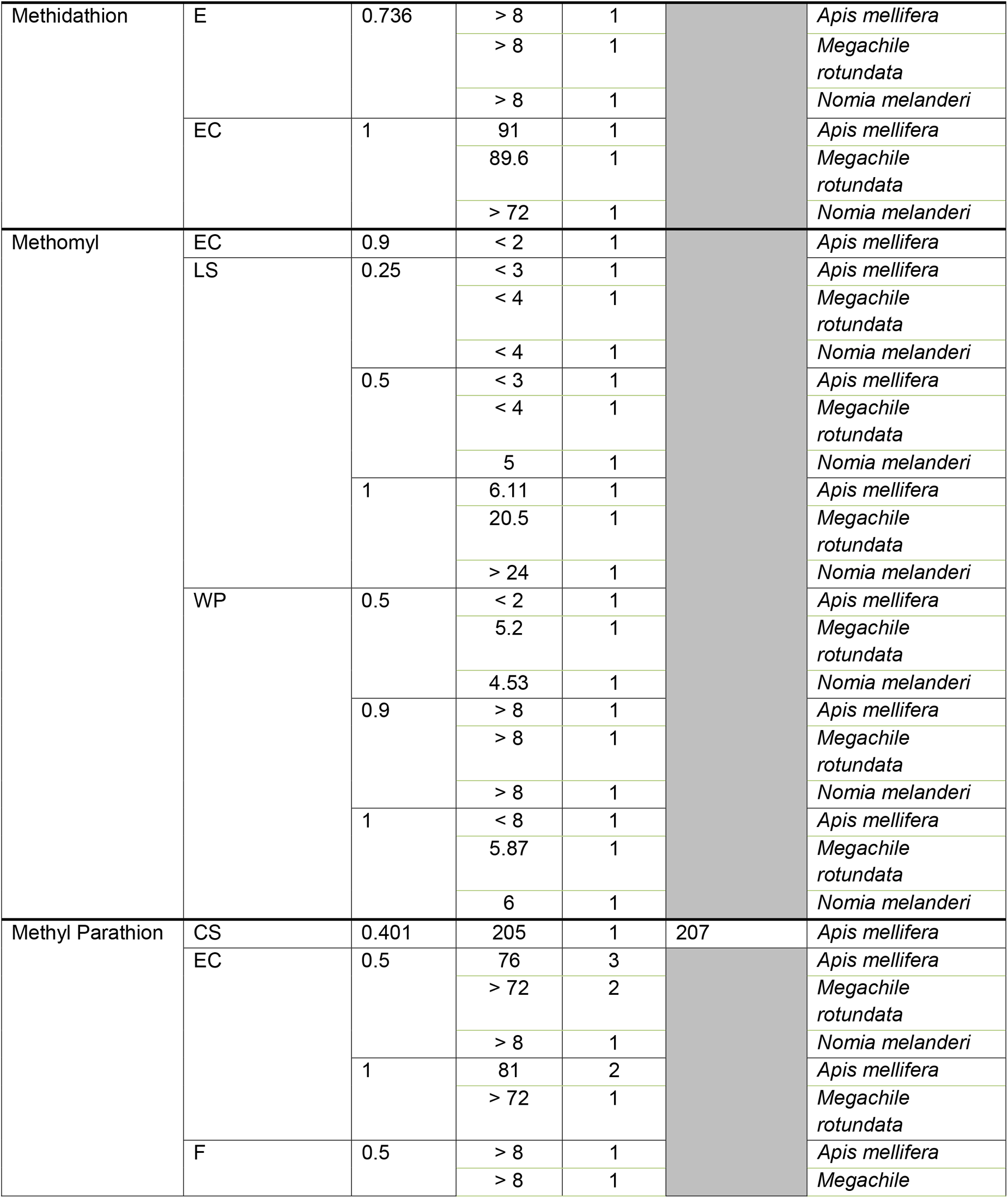
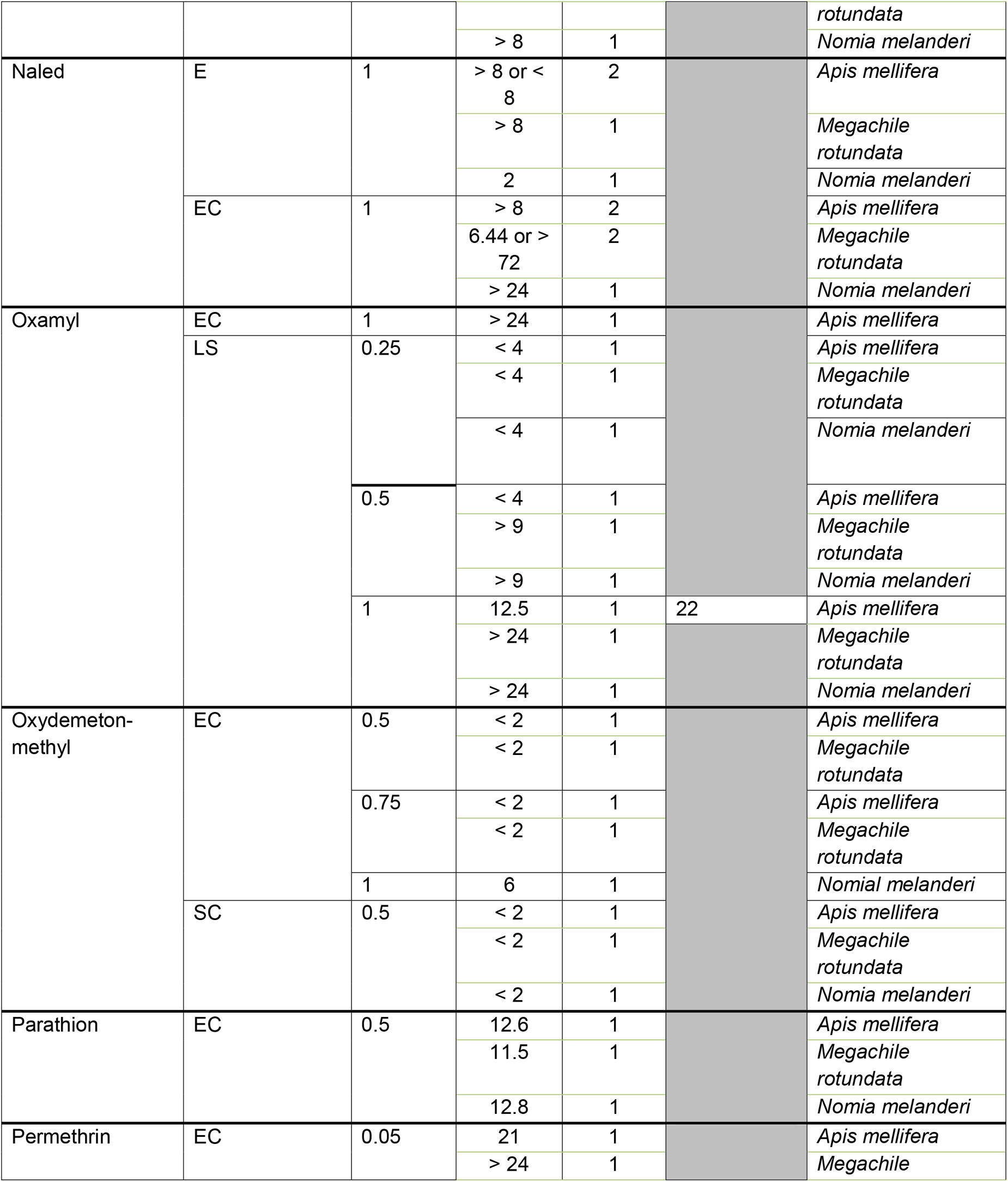
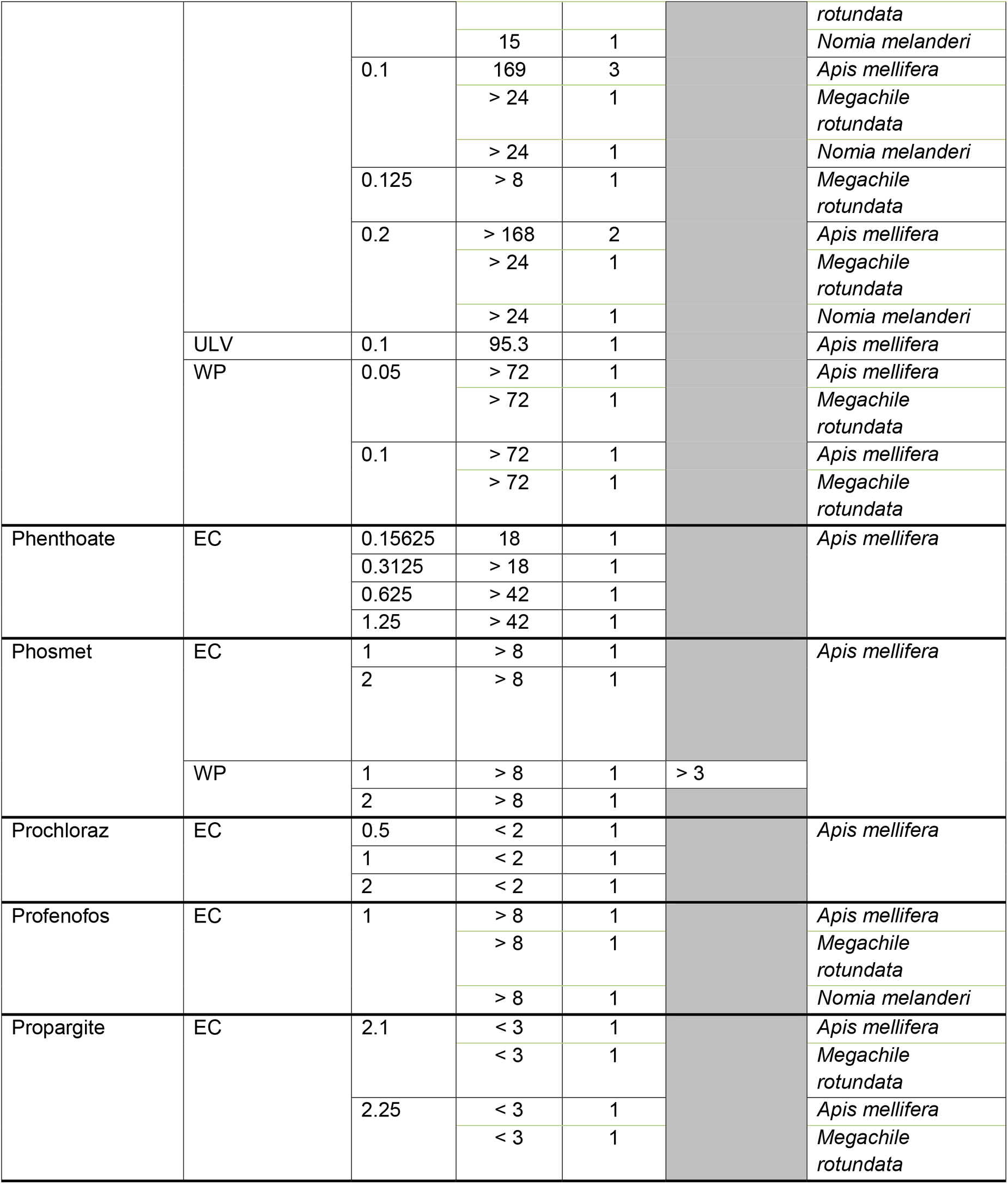
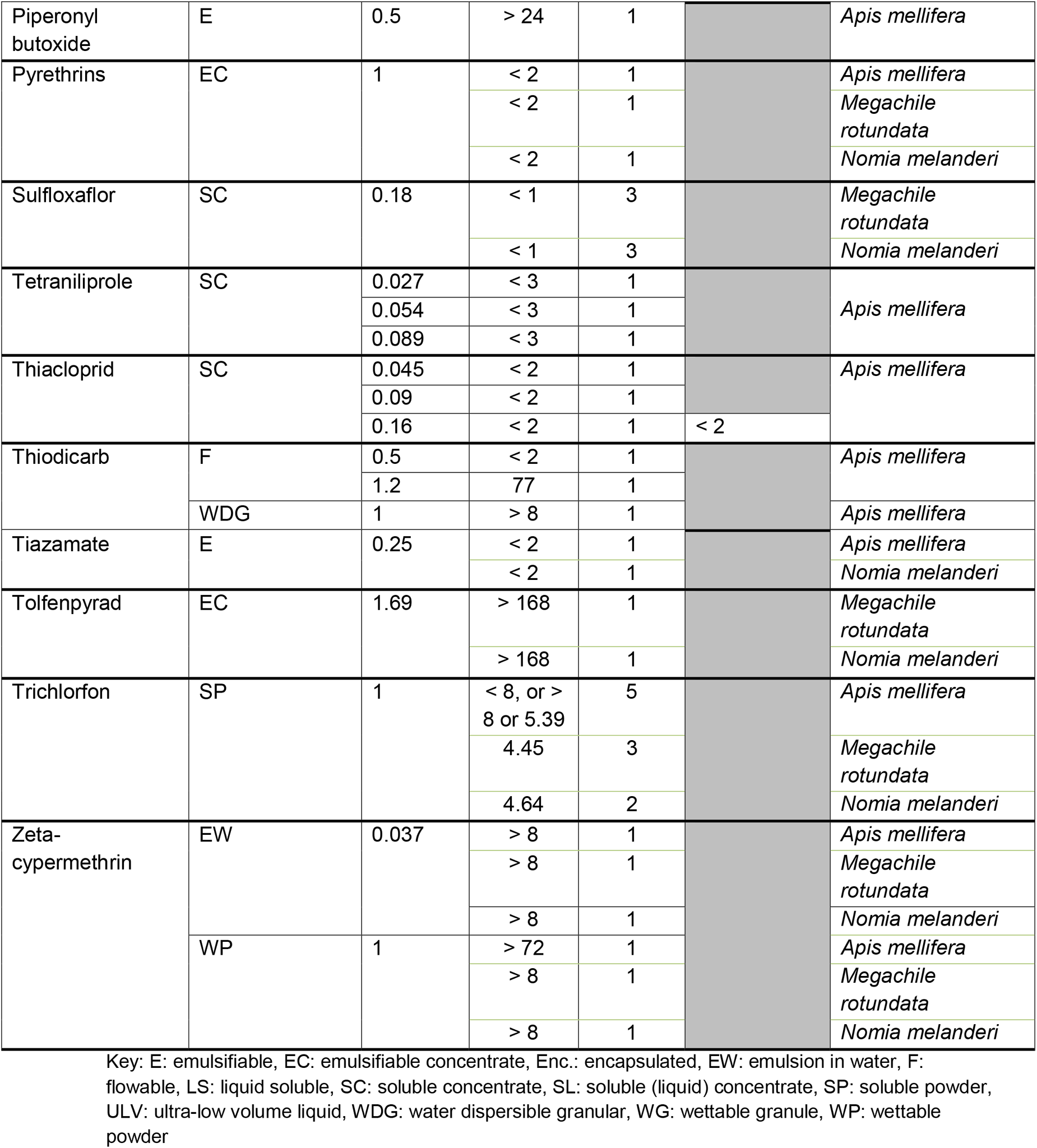
Compiled Calculated RT25 Database

Overall, calculated RT_25_ values from studies matched the extended residual toxicity threshold reported by EPA with a single active ingredient that was not aligned (Figure 3). Notably, this single case, disulfoton emulsifiable concentrate applied at a rate of 1 pound of active ingredient per acre, was close to the extended residual toxicity threshold with a calculated RT_25_ value of 8.86 hours and a published EPA RT_25_ value of 5.5 hours. From our database, we were able to calculate RT_25_ values for an additional 29 active ingredients that were not present in the USEPA’s published database (Figure 4; USEPA, 2014).

**Figure 3:**
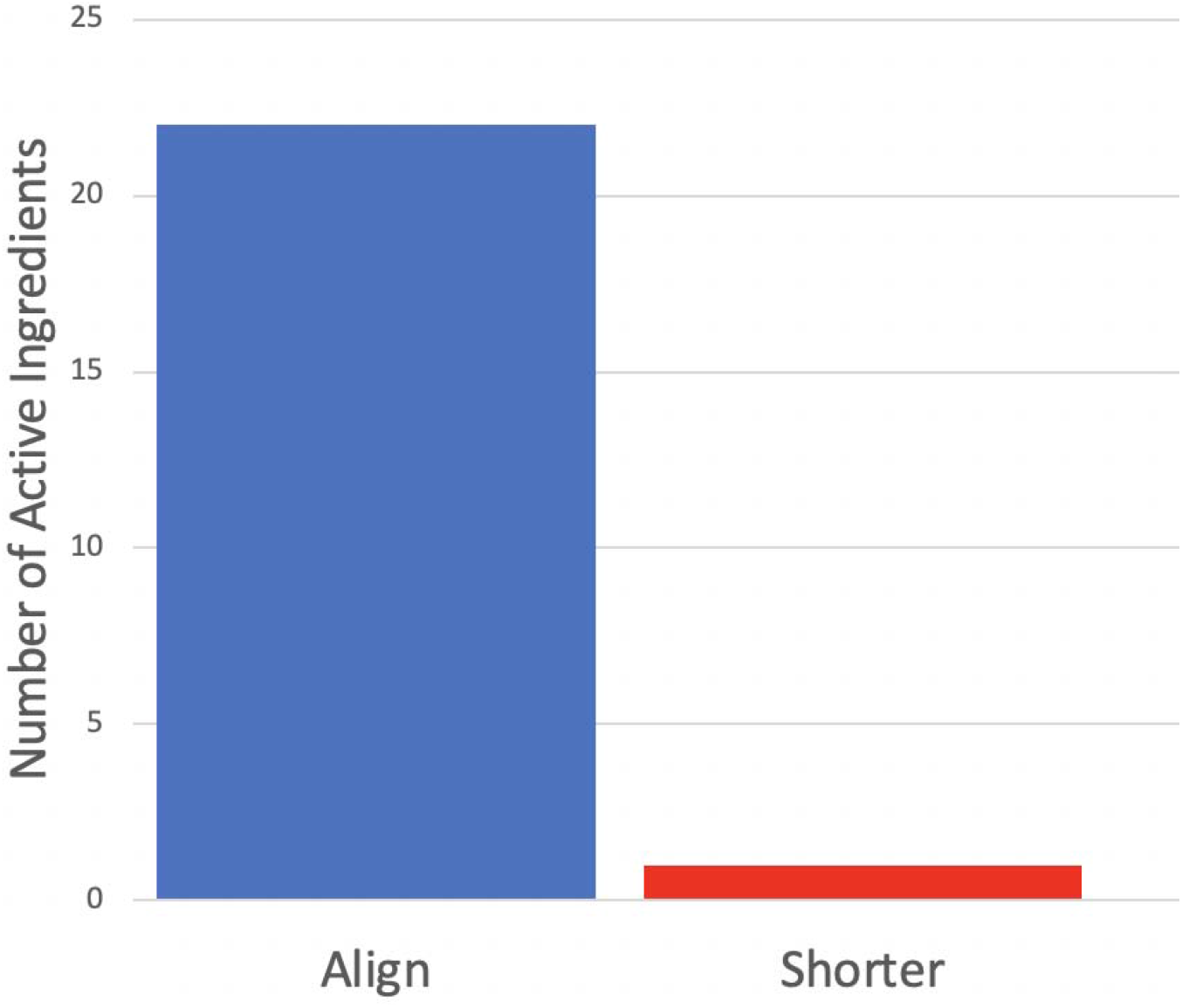
Total number of pesticide active ingredients where RT_25_ values calculated from the literature aligned in their approximations (*e.g*., both reported as greater than 24 hours) or reported the same value with the published by USEPA RT_25_ database (USEPA, 2014) signified by “Aligned”. If the literature and USEPA disagree on the RT_25_ it is depicted with the U reporting a “Shorter” RT_25_ value. We matched formulation and rate when comparing the USEPA values and the literature values.

**Figure 4:**
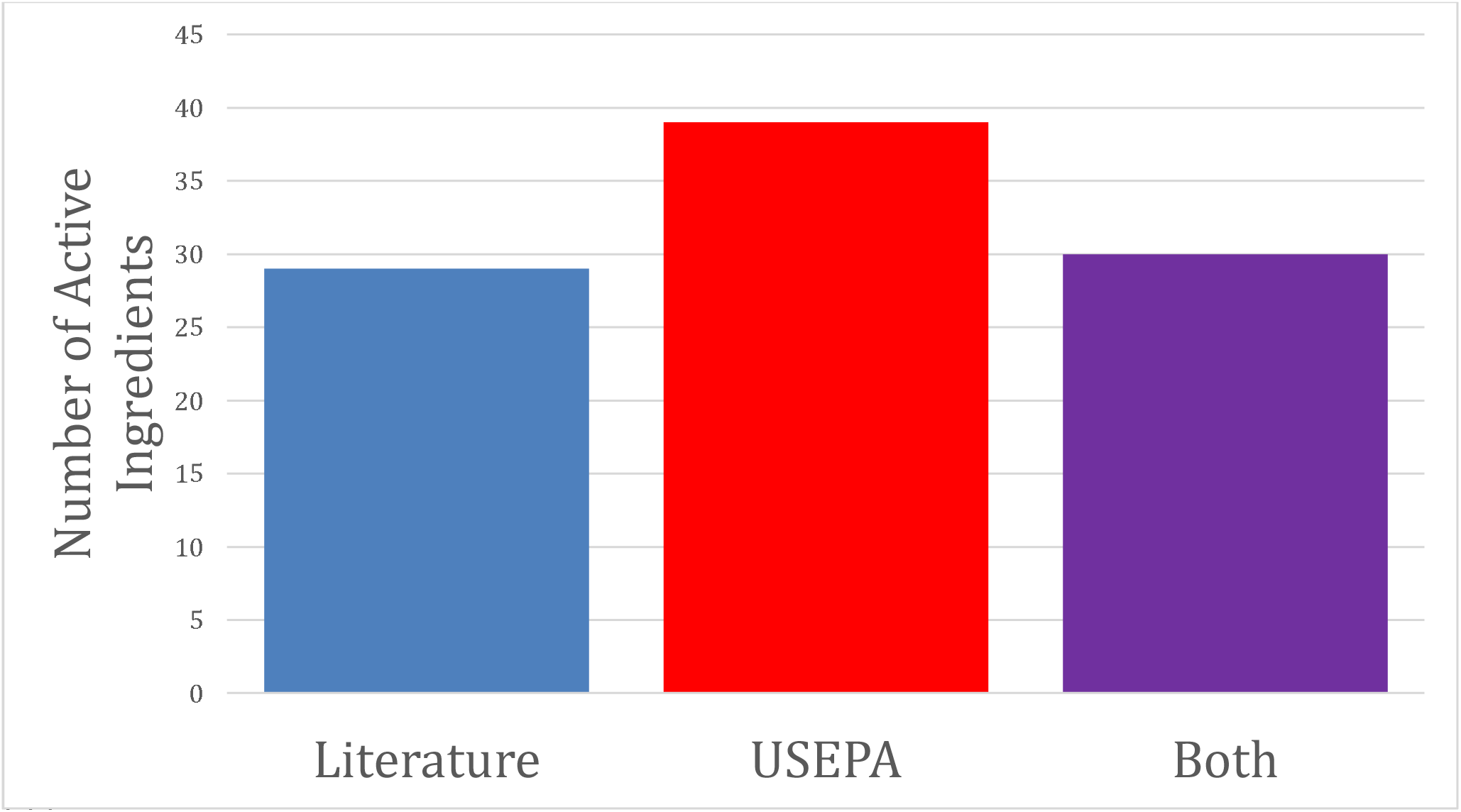
Number of pesticide active ingredients where RT_25_ values could only be calculated from the literature (“Literature”), only from USEPA’s published database (“USEPA”; USEPA, 2014) or where there were RT_25_ values available from both sources (“Both”).

### 3.3 Label language comparisons

Based on the high level of the extended residual toxicity threshold agreement between EPA’s published values (USEPA, 2014) and the database generated through our meta-analysis, we supplemented the EPA database with residual toxicity values for pesticides not previously included. Notably, even after supplementing the EPA database, we were still unable to compare the residual toxicity language on the Environmental Hazards section of labels for one third of pesticides due to a lack of data. Of the remaining labels, a third had residual toxicity warnings that corresponded to RT_25_ values and 27% failed to have any residual toxicity warning despite being toxic to bees. Of the cases where the RT_25_ values did not correspond to residual toxicity statements, 14 of the 22 labels had a statement indicating that the product would remain toxic longer than the RT_25_ value. The other eight had a statement indicating the product would remain toxic shorter than the RT_25_ value (Figure 5).

**Figure 5:**
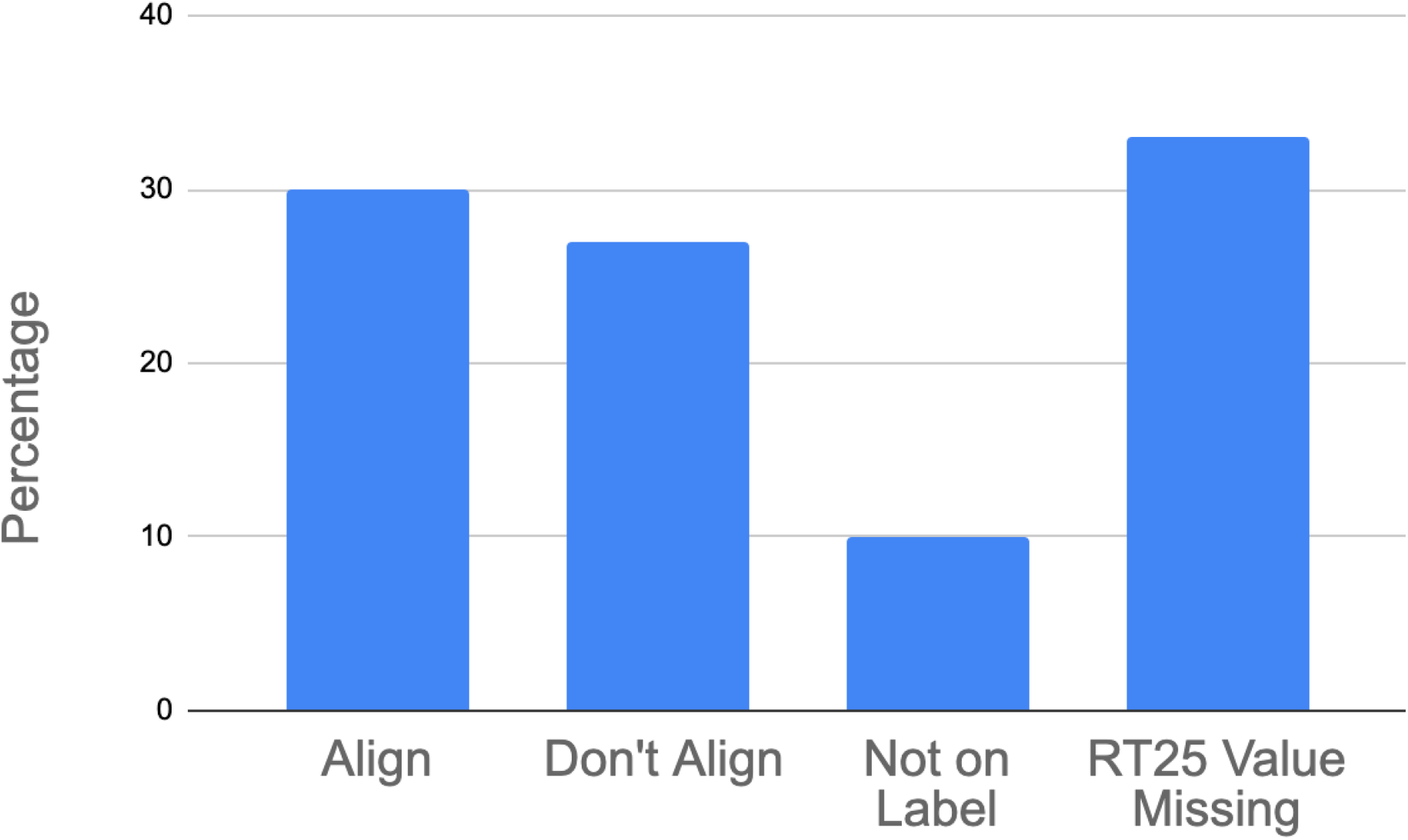
Comparison of pesticide label language indicating residual toxicity in relation to RT_25_ values (calculated from the literature and from USEPA (USEPA, 2014)). Residual toxicity language in the Environmental Hazards section either: (1) aligned with RT_25_ values (“Align”), (2) did not align (“Don’t Align”), (3) lacked residual toxicity language (“Not on Label”) or (4) did not have an RT_25_ value to relate to the label language (“RT_25_ value missing”). Formulation was matched when comparing label language to calculated RT_25_ values.

**Table 3.**
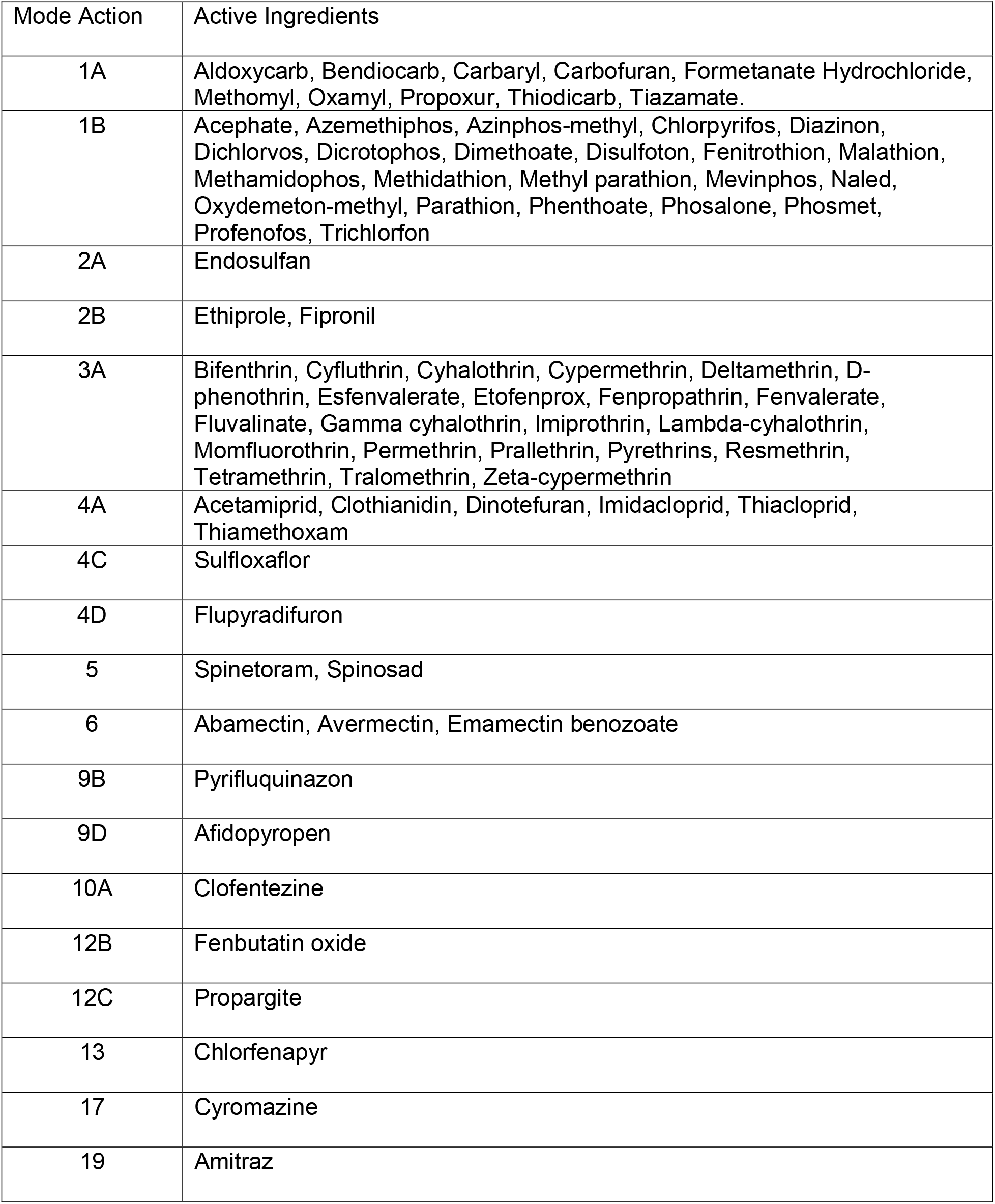
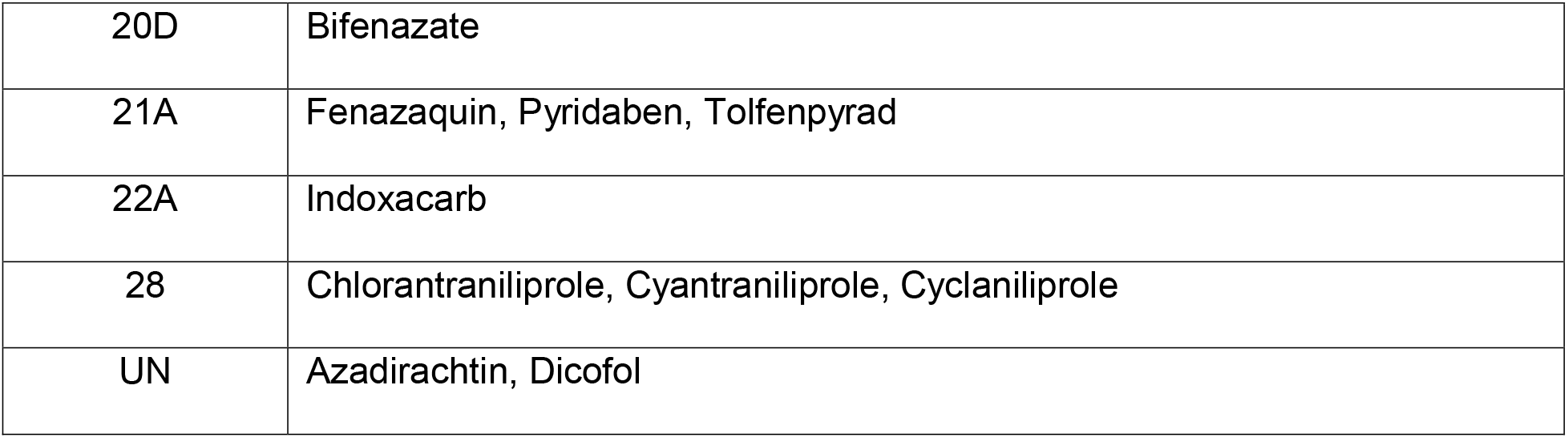
Mode of Action Based on Availability in Insect Resistance Action Committee (2022) for all Active Ingredients in RT_25_ Database (Calculated and USEPA (2014)).

## Discussion

We have developed the most comprehensive database available for RT_25_ values, which greatly expands the publicly accessible values initially published by EPA in 2014. Using published studies, we were able to add 29 active ingredients in addition to the EPA’s values. We demonstrated that while test methodologies varied among published studies, they nonetheless consistent in determining whether a pesticide had extended residual toxicity or not, suggesting that variation in methodologies or the environmental conditions under which the tests were conducted did not result in substantively different conclusions on whether or not an applicator could apply the product at bloom in the evening. Despite our efforts to expand on EPA’s existing RT_25_ database, we found that there remains a paucity of residual toxicity studies. A third of labels in our database did not have corresponding RT_25_ values in either the RT_25_ values calculated as part of our study nor the RT_25_ value estimates published by EPA. We took more variables such as formulation of the active ingredient and application rate into account than may have been necessary when calculating RT_25_ which may have contributed to the lack of comparable RT_25_ values. For example, future studies may choose to not consider variables such as application rate when calculating RT_25_ values to maximize the amount of studies usable for calculating each individual RT_25_ value. Moreover, most studies included in our analysis were published in the 1990s and numerous new active ingredients have since been registered, which illustrates the extent to which researchers have not kept pace with the rate of pesticide product development. Despite these challenges, our meta-analysis expanded EPA’s RT_25_ database and was able to draw attention to widespread misalignment between RT_25_ values and the Pollinator Insect Hazard Statement on pesticide labels that informs pesticide applicators of products with extended residual toxicity.

There was general agreement on whether an active ingredient had extended residual toxicity (i.e., an RT_25_ value >8h) between our meta-analysis and EPA’s database. We found only one deviation across 57 comparable studies of the same active ingredient and application rate. This agreement is remarkable since key aspects of the test methodology were not standardized. Our meta-analysis RT_25_ estimates were often within 1-3 hours of those published by EPA. For example, we calculated the RT_25_ for chlorpyrifos formulated as an emulsifiable concentrate on *A. mellifera* as 17 h compared to the EPA database estimate of 16 h. However, the approach used in this analysis to compare in terms of the extended residual toxicity threshold instead of point estimates may have reduced the influence of methodological variation. Using the extended residual toxicity threshold, the RT_25_ value for the pyrethroid insecticide fenpropathrin at 0.4 lbs ai/A for *A. mellifera* was determined to be greater than 8 h while EPA reported the value as less than 336 hours. Thus, we would deem these two values as the same, because they both support a conclusion of extended residual toxicity, even though the actual estimate of RT_25_ beyond the 8h threshold remains unresolved. Nevertheless, the general agreement between studies on extended residual toxicity is remarkable and suggests that RT_25_ estimates are relatively insensitive to variation in lab technique and weathering conditions.

Our preliminary finding that lab methodology and field weathering conditions are not important sources of variation for RT_25_ should be confirmed experimentally. With respect to lab methodology, we think three factors warrant closer examination, namely the temperature at which the assay is performed, the number of bees held in each test cage and the age of bee used in the test. We report considerable variation in the temperature bees are exposed to in test cage, with temperatures tending to be lower on average compared to EPA guidance. Cooler temperatures could decrease bee activity, leading to less overall contact with the pesticide residue and shorter residual toxicity values (Corbet, 1993). The number of bees in test cages may also influence RT_25_ values by concentrating/diluting the residual pesticide across fewer/greater numbers of bees, resulting in shorter/longer RT_25_ values. We observed that *M. rotundata* and *N. melanderi* had, on average, less bees per cage compared to *A. mellifera* which could lead to less contact per bee to the pesticide residues. Most studies deviated from the age of bees recommended by the EPA, however using less than one day old bees may be distorting, as foraging age bees, which are typically bees that are least three weeks old, are the bees likely to contact weathered residues in the field. Notably, a factor that was largely omitted from most studies was a description of the weathering conditions, such as temperature, humidity, precipitation, and cloud cover. Potentially, weathering conditions may have a larger impact on RT_25_ estimates than variation in laboratory methodology.

We observed trends in RT_25_ values among different rates and formulations of active ingredients. Typically, the higher the application rate of a pesticides, the longer the calculated RT_25_ values. For example, the calculated RT_25_ value for the organophosphate insecticide chlorpyrifos emulsifiable concentrate with *A. mellifera* was 17 hours at the rate of 0.25 lb ai/A and 99 hours RT_25_ time at 0.5 lb ai/A. This suggests that RT_25_ may be different for different application rates, which draws into question the premise of the Pollinating Insect Hazard Statement, where a single residual toxicity statement is meant to cover multiple different use patterns of a pesticide, such as different rates. Notably, new guidance issued by EPA (2017) moves away from relying on the Pollinating Insect Hazard Statement to convey residual toxicity estimates, relying more on specific use directions, where rate and crop are specified. Our results suggest this shift will provide applicators with more guidance on the specific residual times they might experience in the field.

The species of bee used to estimate RT_25_ exhibited notable patterns that should be further investigated. In general, we observed that for the same active ingredient applied at the same rate and formulation *M. rotundata* had longer RT_25_ times compared to *A. mellifera*, and that *N*. *melanderi* had both shorter and longer RT_25_ times compared to *A. mellifera*. Emulsifiable concentrates were associated with the largest difference in RT_25_ estimates among species, with *M. rotundata* consistently having longer RT_25_ values than *A. mellifera* for these formulations. It is unclear what is the source of these patterns. One hypothesis is that *M. rotundata* may be more susceptible to pesticides as this species lacks the ability to detoxify certain synthetic insecticides that are normally metabolized by other bee species (Hayward *et al.,* 2019). Another possible explanation for the difference between bee species could be their size difference. *M. rotundata* has the smallest average size of the three bees we analyzed and, therefore, would have the highest ratio of surface area to body volume. The higher surface area to body volume ratio would expose the bees to higher levels of contact exposure, and longer RT_25_ times than *A. mellifera* (Wisk *et al.,* 2014). LD_50_ values for each species could also contribute to the differences. Bee species, specifically *M. rotundata* and *N. melanderi*, also tend to vary in LD_50_ values for many active ingredients such as dicrotophos, malathion, and fipronil compared to *A. mellifera* (Devillers and Pham-Delegue, 2002). Furthermore, in another study, specifically for the organophosphate cyhalothrin, *M. rotundata* yielded the highest LD_50_ values, *A. mellifera* yielded intermediate values, and *N. melanderi* yielded lowest values (Mayer *et al.,* 1998). Little research has been done into the effects of differing formulations on the residual toxicity across bee species. A species comparative study would be useful to determine what variables (e.g., differences in behavior, different physiology, etc.) contribute to the differing residual toxicity values. Currently, EPA publicly reports (EPA, 2014) RT_25_ times primarily for *A. mellifera*, with limited data available on other species of pollinating bees. Differences in species residual toxicity times have been noticed in the past (Johansen *et al*., 1983; Mayer *et al*., 1997) but there have been no in-depth studies designed to comparatively characterize RT_25_ estimates for different species, let alone resolve the mechanisms by which bees may respond to the dissipation of residues differently. Certainly, variation in RT_25_ estimates for different bee species would be important information for pesticide applicators, particularly if they are using residual times for bee species with the shortest RT_25_ values.

The finding from our study that is of greatest concern to pesticide applicators was widespread misalignment between RT_25_ values and statements of residual toxicity in the Pollinating Insect Hazard Statement. Of the pesticide labels we were able to compare to calculated RT_25_ values, almost a third were inaccurate in the wording of their Pollinating Insect Hazard Statement. For example, the formulated end-use product Perm-Up 3.2 EC (USEPA registration number 70506-9) containing the pyrethroid insecticide permethrin indicates the product should not be applied while bees are “actively visiting” suggesting a less than 8-hour residual toxicity time. However, the residual toxicity studies for permethrin consistently indicated RT_25_ values greater than 8 hours even at the lowest application rate calculated, 0.05 lb ai/A. Although this finding is concerning, some of these discrepancies may arise from our assumption that all pesticides with the same active ingredient and applied at the same rate have similar RT_25_ values. Potentially, pesticide products may have different residual times owing to features independent of the active ingredient, such as inert ingredients. Our assumption that RT_25_ can be generalized across products containing the same active ingredient is supported by our findings that RT_25_ estimates were largely consistent for active ingredients across studies and relative to estimates published by EPA (EPA, 2014). Nevertheless, we suggest caution in interpreting our results since the number of different products used to estimate RT_25_ values for each active ingredient tended to be dwarfed by the total number of registered products containing those ingredients on the market. Regardless, our study indicates that either there is high variability in residual toxicity between pesticides containing the same active ingredients, which calls into the question efforts like USEPA’s to publish RT_25_ values based on active ingredients, or the Pollinating Insect Hazard Statement on existing pesticide labels aligns poorly with RT_25_ values. Our data currently suggests the latter problem predominates, resulting in pesticide applicators lacking a reliable piece of information to mitigate exposure to bees during bloom.

One thing is clear from our study; there remain large gaps in our database of RT_25_ estimates. Although this database is the most comprehensive to date, and expands on published values by EPA, the lack of publicly accessible RT_25_ estimates is something we hope researchers will make a concerted effort to address. We also encourage EPA to review its existing data from registrants, which is unavailable to researchers, pesticide applicators and the public, and fill gaps in its public-facing database. Alternatively, EPA could develop a mechanism to release registrant-collected residual toxicity data to the public to enable researchers to develop such a database independently. While estimating residual toxicity has been a part of the pesticide risk assessment process for decades, its relevance continues with new guidance around label language that foregrounds RT_25_ values beyond the Environmental Hazard section to the crop-specific directions for use on the label (EPA, 2017). The need to create a basis for evaluation of these changes is not only important for pesticide applicators who are seeking instruction to protect bees from exposure, but for the sustainable management of managed bee stocks and wild bee communities.

## Conclusions

Concerns over the communication of the Pollinating Insects Hazards Statement on pesticide labels have arisen in the past. Through our efforts, we were successfully able to create a compendium of RT_25_ values that could be used to determine if pesticide label language aligns with calculated active ingredient RT_25_ values. There was noticeable variation in species and application rate which could call into question whether a single Pollinating Insects Hazards Statement is sufficient to fully communicate the hazards of a pesticide product. Further comparison of the calculated values to published EPA values revealed that lab methodology does not seem to affect RT_25_ values as seen from comparison of study values to EPA, though field conditions during the weathering of the pesticide may need to be explored further. Comparing a combined database of published EPA values and our calculated RT_25_ values to label language showed significant misalignment in Pollinating Insect Hazard Statements. The variation in residual toxicity remains an emerging field of research that must be addressed to ensure the applications of pesticides is occurring in a safe manner to minimize the risk towards pollinating bees.

## Supporting information

RT25 Calculation Code

All Residual Toxicity Data

PRISMA Flow Diagram

## Acknowledgements

We thank Drs. Daniel Schmehl and Allen Olmstead at Bayer CropScience for unpublished residual toxicity data and technical assistance with calculations of RT_25_ as well as Dr. Theresa Pitts-Singer for compiling residual toxicity studies from the Western Alfalfa Seed Growers Association. We also greatly thank Matthew T. Bucy for compiling a pesticide label database and producing preliminary data from residual toxicity studies. Finally, we owe a debt of gratitude to Dr. Thomas Steeger for reviewing an advanced draft of this manuscript. The research was funded from a grant the Western IPM Center and from the Western Sustainable Agriculture and Education Research and Education Grant.

## References

Akca I, Tuncer C, Guler A, Sarugan I. 2009. Residual toxicity of 8 different insecticides on honey bee (*Apis mellifera* Hymenoptera: Apidae). Journal of Animal and Veterinary Advances 8:436–440.

Bailey J, Scott-Dupree C, Harris R, Tolman J, Harris B. 2005. Contact and oral toxicity to honey bees (*Apis mellifera*) of agents registered for use for sweet corn insect control in Ontario, Canada. EDP Sciences 36:623–633. DOI: 10.1051/apido:2005048.

Barmaz S, Potts SG, Vighi M. 2010. A novel method for assessing risks to pollinators from plant protection products using honeybees as a model species. Ecotoxicology 19:1347–1359. DOI: 10.1007/s10646-010-0521-0.

Botías C, David A, Hill EM, Goulson D. 2017. Quantifying exposure of wild bumblebees to mixtures of agrochemicals in agricultural and urban landscapes. Environmental Pollution 222:73–82. DOI: 10.1016/j.envpol.2017.01.001.

Bucy M, Melathopoulos A. 2020. Labels of insecticides to which Oregon honey bee (*Apis mellifera* L.) hives could be exposed do not align with federal recommendations in their communication of acute and residual toxicity to honey bees. Pest Management Science 76:1664–1672. DOI: 10.1002/ps.5685.

Chauzat M-P, Martel A-C, Blanchard P, Clément M-C, Schurr F, Lair C, Ribière M, Wallner K, Rosenkranz P, Faucon J-P. 2010. A case report of a honey bee colony poisoning incident in France. Journal of Apicultural Research 49:113–115. DOI: 10.3896/IBRA.1.49.1.22.

Clinch P. 1967. The residual contact toxicity to honey bees of insecticides sprayed on to white clover (*Trifolium repens* I.) in the laboratory. New Zealand Journal of Agricultural Research 10:289–300. DOI: 10.1080/00288233.1967.10425136.

Corbet SA, Fussell M, Ake R, Fraser A, Gunson C, Savage A, Smith K. 1993. Temperature and the pollinating activity of social bees. Ecological Entomology 18:17–30. DOI: 10.1111/j.1365-2311.1993.tb01075.x.

Devillers J, Pham-Delegue M-H. 2002. Honey bees: estimating the environmental impact of chemicals. New York: Taylor & Francis, 112.

Fischer D, Moriarty T. 2011. Overview of honey bee biology, In: Pesticide risk assessment for pollinators: summary of a SETAC pellston workshop. Washington: SETAC Press, 8.

Hayward A, Beadle K, Singh KS, Exeler N, Zaworra M, Almanza M-T, Nikolakis A, Garside C, Glaubitz J, Bass C, Nauen R. 2019. The leafcutter bee, *Megachile rotundata*, is more sensitive to N-cyanoamidine neonicotinoid and butanolide insecticides than other managed bees. Nature Ecology & Evolution 3:1521–1524. DOI: 10.1038/s41559-019-1011-2.

Hooven L, Sagili R, Johansen E. 2013. How to reduce bee poisoning from pesticides. Corvallis: Oregon State University Extension Publications.

Johansen C. 1972. Toxicity of field-weathered insecticide residues to four kinds of bees. Environmental Entomology 1:393–394.

Johansen C, Baird C. 1972. Small-scale bee poisoning tests with honey bees (HB), alkali bees (AB), and alfalfa leafcutting bees (LB). In: Bee research investigations. Pullman: Washington State University, 2–3, 13–17.

Johansen C, Eves J. 1971. Small-scale bee poisoning tests with honey bees (HB), alkali bees (AB), alfalfa leafcutter bees (LB), and bumble bees (BB). In: Bee research investigations. Pullman: Washington State University, 2–3, 13–17.

Johansen C, Kious C, George Schultz, Gupta R, Madsen R, Robinson W. 1977. Investigation of the bee poisoning hazards of microencapsulated methyl parathion (penncap-M). In: Bee research investigations. Pullman: Washington State University, 1–5,16,19–20.

Johansen C, Kious C, Mayer D. 1981. Small-scale poisoning tests with honey bees, alkali bees, and alfalfa leafcutting bees. In: Bee research investigations. Pullman: Washington State University, 1–3.

Johansen C, Mayer D, Baird C. 1973. Small-scale bee poisoning tests with honey bees (HB), alkali bees (AB), and alfalfa leafcutting bees (LB). In: Bee research investigations. Pullman: Washington State University, 3–4, 9–14.

Johansen C, Mayer D, Eves J, Kious C. 1983. Pesticides and bees. Environmental Entomology 12:1513–1518.

Johansen C, Mayer D, Kious C. 1984. Small-scale poisoning tests with honey bees and alfalfa leafcutting bees. In: Bee research investigations. Pullman: Washington State University, 1– 2.

Johansen C, Mayer D, Kious C, Sheffield C. 1983. Small-scale poisoning tests with honey bees, alkali bees, and alfalfa leafcutting bees. In: Bee research investigations. Pullman: Washington State University, 1.

Johansen C, Mayer D, Madsen R, Robinson W. 1975. Small-scale bee poisoning tests with honey bees (HB), alkali bees (AB), and alfalfa leafcutting bees (LB). In: Bee research investigations. Pullman: Washington State University, 1–2, 12–15.

Johansen C, Mayer D, Robinson W, Gupta R, Spann J, Madsen R. 1976. Small-scale bee poisoning tests with honey bees (HB), alkali bees (AB), and alfalfa leafcutting bees (LB). In: Bee research investigations. Pullman: Washington State University,1–2,13–16.

Keshlaf M, Basta A, Spooner-Hart R. 2013. Assessment of toxicity of fipronil and its residues to honey bees. Mellifera 13:30–38.

Kiljanek T, Niewiadowska A, Gaweł M, Semeniuk S, Borzęcka M, Posyniak A, Pohorecka K. 2017. Multiple pesticide residues in live and poisoned honeybees – Preliminary exposure assessment. Chemosphere 175:36–44. DOI: 10.1016/j.chemosphere.2017.02.028.

Kim B-S, Park Y-K, Lee Y-H, Joeng M-H, You A-S, Yang Y-J, Kim J-B, Kwon O-K, Ahn Y-J. 2008. Honeybee acute and residual toxicity of pesticides registered for strawberry. The Korean Journal of Pesticide Science 12:229–235.

Kious C, Schultz G, Johansen C. 1979. Small-scale bee poisoning tests with honey bees (*Apis mellifera*). In: Bee research investigations. Pullman: Washington State University, 3–4.

Mayer D. 2001. Residual bee poisoning bioassay. In: Integrated pest and pollinator investigations. Prosser: Washington State University, 1–2.

Mayer DF, Johansen C. 1985. Pollinator protection and Acephate (Orthene) insecticide. In: Agricultural Research. 125:207–210.

Mayer D, Johansen C, Shanks C, Lunden J. Insecticide residues. In: Methomyl and honey bees. Prosser: Washington State University, 46–47.

Mayer D, Johansen C, Shanks C, Pike K. 1987. Effects of Fenvalerate Insecticide on Pollinators. Journal of Entomological Society of British Columbia 84:39–45.

Mayer D, Kovacs G, Brett B, Bisabri B. 2001. The effects of spinosad insecticide to adults of *Apis mellifera*, *Megachile rotundata* and *Nomia melanderi* (Hymenoptera: Apidae). International Journal of Horticultural Science 7:93–97.

Mayer D, Kovacs G, Lunden J. 1998. Field and laboratory tests on the effects of cyhalothrin on adults of *Apis mellifera, Megachile rotundata* and *Nomia melanderi*. Journal of Apicultural Research 37:33–37. DOI: 10.1080/00218839.1998.11100952.

Mayer D, Lunden J. 1999. Field and laboratory tests of the effects of fipronil on adult female bees of *Apis mellifera*, *Megachile rotundata*, and *Nomia melanderi*. Journal of Apicultural Research 38:191–197.

Mayer D, Lunden J. 1999. Residual bee poisoning bioassay. In: Integrated pest and pollinator investigations. Prosser: Washington State University, 2–5.

Mayer D, Lunden J, Husfloen M. 1991. Residual bee poisoning bioassay. In: Integrated pest and pollinator investigations. Prosser: Washington State University, 1–2.

Mayer D, Lunden J, Jasso M. 1996. Residual bee poisoning bioassay. In: Integrated pest and pollinator investigations. Prosser: Washington State University, 1–4.

Mayer D, Lunden J, Jasso M. 1997. Residual bee poisoning bioassay. In: Integrated pest and pollinator investigations. Prosser: Washington State University, 1–4.

Mayer D, Lunden J, Johansen C. 1985. Small scale bee poisoning bioassay. In: Bee research investigations. Prosser: Washington State University, 2–5.

Mayer D, Lunden J, Kovacs G. 1997. Susceptibility of four bee species (Hymenoptera: Apoidea) to field weathered insecticide residues. Journal of Entomological Society of British Columbia 94:27–30.

Mayer D, Lunden J, Miliczky E. 1988. Residual bee poisoning bioassay. In: Integrated pest and pollinator investigations. Prosser: Washington State University, 1–5.

Mayer D, Lunden L, Rathbone L, Johansen C. 1986. Residual bee poisoning bioassays. In: Bee research investigations. Prosser: Washington State University, 2–6.

Mayer D, Lunden J, Rathbone L, Miliczky E, Johansen CA. 1987. Residual bee poisoning bioassay. In: Bee research investigations. Prosser: Washington State University, 1–5.

Mayer D, Patten K, Macfarlane R, Shanks C. 1994. Differences between susceptibility of four pollinator species (Hymenoptera:Apoidea) to field weathered insecticide residues. Melanderia 50.

Mayes M, Thompson G, Husband B, Miles M. 2003. Spinosad toxicity to pollinators and associated risk. Reviews of Environmental Contamination and Toxicology 179:37–71.

Mode of Action Classification | Insecticide Resistance Management. Available at https://irac-online.org/mode-of-action/classification-online/ (accessed March 1, 2023).

Office of Chemical Safety and Pollution Prevention. 2012. White paper in support of the proposed risk assessment process for bees chapter 4: characterization of ecological effects.

Pashte V, Patil C. 2017. Evaluation of persistence of insecticide toxicity in honey bees (Apis mellifera L.). Indian Journal of Biochemistry and Biophysics 54:150–155.

R Core Team. 2021. R: A language and environment for statistical computing. R Foundation for Statistical Computing.

Sanchez-Bayo F, Goka K. 2014. Pesticide residues and bees – a risk assessment. PLoS ONE 9:e94482. DOI: 10.1371/journal.pone.0094482.

Smodiš Škerl MI, Velikonja Bolta Š, Baša Česnik H, Gregorc A. 2009. Residues of pesticides in honeybee (Apis mellifera carnica) bee bread and in pollen loads from treated apple orchards. Bulletin of Environmental Contamination and Toxicology 83:374–377. DOI: 10.1007/s00128-009-9762-0.

Summit Agro, Harvanta 50SL INSECTICIDE.

The Honey Bee Health Coalition. 2019. Best management practices for hive health a guide for beekeepers.

Tosi S, Costa C, Vesco U, Quaglia G, Guido G. 2018. A 3-year survey of Italian honey bee-collected pollen reveals widespread contamination by agricultural pesticides. The Science of the Total Environment 615:208–218. DOI: 10.1016/j.scitotenv.2017.09.226.

United States Environmental Protection Agency. 2016. Guidance on exposure and effects testing for assessing risks to bees.

United States Environmental Protection Agency. 2012. Honey bee toxicity of residues on foliage.

United States Environmental Protection Agency. 2012. Label review manual chapter 8: environmental hazards.

United States Environmental Protection Agency. 2014. Residual time to 25% bee mortality (RT25) data. Available at https://www.epa.gov/pollinator-protection/residual-time-25-bee-mortality-rt25-data (accessed February 21, 2023).

United States Protection Agency. 2017. U.S. Environmental protection agency’s policy to mitigate the acute risk to bees from pesticide products chapter 5: modifications to the environmental hazards section of pesticide labels.

Vinchesi A, Boyle N, Walsh D. 2013. Studies on alkali bees and pollinator pesticide safety in Washington State.

Waller G, Estesen B, Buck N, Taylor K, Crowder L. 1988. Residual life and toxicity to honey bees (Hymenoptera:Apidae) of selected pyrethroid formulations applied to cotton in arizona. Journal of Economic Entomology 81:1022–1026. DOI: 10.1093/jee/81.4.1022.

Walsh D. 2010. Insecticide Efficacy Trials 2008-2009. In: Integrated pest management on alfalfa seed: a two-year report 2008-2009. Las Vegas: Western Alfalfa Seed Growers Association, 1–7.

Walsh D, Boydston R, O’Neal S. 2008. 2005-2007 Alfalfa seed research report.

Walsh D, Waters T, O’Neal S, Groenendale D, Peng W, Vinchesi A, Piraneo T. 2011. Pest and pollinator management on alfalfa seed 2011.

Walsh D, Wine E, Groenendale D, Vinchesi A, Boyle N. 2016. Pest and pollinator management on alfalfa seed 2015.

Wickham H, Averick M, Bryan J, Chang W, McGowan LD, François R, Grolemund G, Hayes A, Henry L, Hester J, Kuhn M, Pedersen TL, Miller E, Bache SM, Müller K, Ooms J, Robinson D, Seidel DP, Spinu V, Takahashi K, Vaughan D, Wilke C, Woo K, Yutani H. 2019. Welcome to the tidyverse. Journal of Open Source Software, 4:43:1686. doi:10.21105/joss.01686.

Wiese I. The susceptibility of honeybees to some insecticide spray formulations used on citrus. South African Journal of Agricultural Science 5:557–588.

Winston ML. 1991. The Biology of the Honey Bee: Cambridge, MA: Harvard University Press.

Wisk J, Pistorius J, Beevers M, Bireley R, Browning Z, Chauzat M, Nikolakis A, Overmyer J, Rose R, Sebastien R, Vaissière B, Maynard G, Kasina M, Nocelli R, Scott-Dupree C, Johansen E, Brittain C, Coulson M, Dinter A, Vaughan M. 2014. Assessing exposure of pesticides to bees. Pesticide Risk Assessment for Pollinators. John Wiley & Sons, Ltd, 45–74. DOI: 10.1002/9781118852408.ch7.

