## Supplementary material for "Systematic review of residual toxicity studies of pesticides to bees and comparison to language on pesticide labels using data from studies and the Environmental Protection Agency": RT25 Calculation Code

```
######################################################################
####################################
##
##              Calculating RT25 values
##
## Project: RT Literature Review
## Author: Leah Swanson
## Date Created: July 13, 2022 by Allen Olmstead
## Last edited: July 27, 2022 by Leah Swanson
##
## Input:
##    RT_data: csv of residual toxicity data
##
## Outputs:
##    A logistic regression and RT25 value
######################################################################
####################################

######################################################################
####################################
#
#
### EACH TIME YOU MAKE AN EDIT: Update "Last edited" field in header
and the name of the editor" #########
#
###
######################################################################
####################################

######################################################################
####################################
### BEFORE YOU RUN THIS SCRIPT
### 1. This script pulls in a csv version of the RT literature review
spreadsheet. MAKE SURE
###    IT IS UP TO DATE. SAVE THE MOST RECENT RT LITERATURE REVIEW
SPREADSHEET AS THE XLSX FILE, JUST
###    TO BE SAFE
### 2. Set working directory
###############################


library(tidyverse)

# read data set in
setwd("/Users/leahswanson/Desktop")
dat <- read_csv("Nomia Johansen et al. 1975 imdacloprid.csv",
show_col_types = FALSE)

##Getting number of mortality and survived
##create a function to calculate be mortalities
```

```
mortality <- function(x) {
  deaths <- (dat$'Mortality (%)'/100) * dat$'number_of_bees'
  return(deaths)
}
##Create a function to calculate alive bees
survival <- function(x2) {
  alive <- dat$'number_of_bees' - ((dat$'Mortality (%)'/100) *
dat$'number_of_bees')
  return(alive)
}
##Apply the functions to the dataframe and create a new column to hold
the values
dat$mort <- apply(X = dat, MARGIN = 1, FUN = mortality)
dat$surv <- apply(X = dat, MARGIN = 1, FUN = survival)

##round values to the nearest whole number
dat$mort <- round(dat$mort, digits = 0)
dat$surv <- round(dat$surv, digits = 0)

######################################################################
#####
# fit a binomial glm (logistic regression)
bin_fit <- glm(cbind(mort, surv) ~ time, dat, family = binomial)

# vectors for graphing model fit
x <- seq(0, 80, length = 1000)
y_fit <- predict(bin_fit, data.frame(time = x), type = "response",
                 se.fit = TRUE)
lcl <- y_fit$fit - 1.96*y_fit$se.fit      # 1.96 = 95% CI (Wald)
ucl <- y_fit$fit + 1.96*y_fit$se.fit

# plot binomial glm fit
ggplot() +
  geom_ribbon(data = NULL, aes(x = x, ymin = lcl, ymax = ucl),
              color = NA, alpha = 0.2) +
  geom_line(data = NULL, aes(x, y_fit$fit), color = "blue", size = 1)
+
  geom_point(data = dat, aes(time, mort/dat$number_of_bees), color =
"red", size = 2) +
  theme_bw() +
  labs(x = "Time (hr)", y = "Mortality (proportion)")

# test for overdisperion using DHARMa package function
DHARMa::testDispersion(bin_fit)

######################################################################
#####
# fit quasibinomial to account for overdispersion
qbin_fit <- glm(cbind(mort, surv) ~ time, dat, family = quasibinomial)
```

```r
summary(qbin_fit)
# vectors for graphing model fit
y_qfit <- predict(qbin_fit, data.frame(time = x), type = "response",
                  se.fit = TRUE)
qlcl <- y_qfit$fit - 1.96*y_qfit$se.fit
qucl <- y_qfit$fit + 1.96*y_qfit$se.fit

# fit a quasibinomial glm fit
ggplot() +
  geom_ribbon(data = NULL, aes(x = x, ymin = qlcl, ymax = qucl),
              color = NA, alpha = 0.2) +
  geom_line(data = NULL, aes(x, y_qfit$fit), color = "blue", size = 1)
+
  geom_point(data = dat, aes(time, mort/30), color = "red", size = 2)
+
  geom_hline(yintercept = 0.25, lty = 2) +
  geom_vline(xintercept = 17, lty = 2) +   # value from section below
  theme_bw() +
  labs(x = "Time (hr)", y = "Mortality (proportion)")

######################################################################
#####
# starting values for algorithm
start_low <- 6.5   # pick a value below that estimated from graph
start_high <- 7   # pick a value above that estimated from graph
target <- 0.25        # the proportional mortality you want a time for

# wrapper function for predict
get_mort <- function(x) {
  predict(bin_fit, data.frame(time = x), type = "response")
}

# initialize a tibble to populate
optim_time <- tibble(lo = start_low, hi = start_high,
                     lo_pred = get_mort(start_low),
                     hi_pred = get_mort(start_high))

# loop for algorithm that fills out tibble with values
for(i in 1:99) {
  diff <- optim_time[i, 2] - optim_time[i, 1]
  if(optim_time$lo_pred[i] - target < target - optim_time$hi_pred[i])
{
    optim_time[i + 1, 1] <- optim_time[i, 1]
    optim_time[i + 1, 2] <- optim_time[i, 2] - diff/2
  } else {
    optim_time[i + 1, 1] <- optim_time[i, 1] + diff/2
    optim_time[i + 1, 2] <- optim_time[i, 2]
  }
  optim_time[i + 1, 3] <- get_mort(optim_time$lo[i + 1])
  optim_time[i + 1, 4] <- get_mort(optim_time$hi[i + 1])
```

```
}

# check the last rows to verify sufficient convergence
tail(optim_time)

######################################################################
#####
# nls fits for logistic equation
dat <- mutate(dat, mort2 = mort/(mort + surv))

# 2 parameter logistic
nls_fit2 <- nls(mort/(mort + surv) ~ 1/(1 + exp(-k*(time - x0))),
                dat, start = list(k = -0.1, x0 = 150))
# 3 parameter logistic
nls_fit3a <- nls(mort/(mort + surv) ~ max/(1 + exp(-k*(time - x0))),
                 dat, start = list(max = 1, k = -0.1, x0 = 150))
# alternative 3 parameter logistic
nls_fit3b <- nls(mort/(mort + surv) ~ (1 - min)/(1 + exp(-k*(time -
x0))) + min,
                 dat, start = list(k = -0.1, x0 = 150, min = 0))
# 4 parameter logistic
nls_fit4 <- nls(mort/(mort + surv) ~ (max - min)/(1 + exp(-k*(time -
x0))) + min,
                dat, start = list(max = 0.8, k = -0.1, x0 = 150, min =
0))

# tibble for fit values for graphing
fitz <- tibble(fit_x = rep(x, 3),
               fit_y = c(predict(nls_fit2, data.frame(time = x)),
                         predict(nls_fit3b, data.frame(time = x)),
                         predict(nls_fit4, data.frame(time = x))),
               Fit = rep(c("2", "3b", "4"), each = length(x)))

# graph the data with all nls fits
ggplot() +
  geom_line(data = fitz, aes(fit_x, fit_y, color = fitz$Fit), size =
1) +
  geom_point(data = dat, aes(time, mort2), color = "red", size = 2) +
  theme_bw() +
  labs(x = "Time (hr)", y = "Mortality (proportion)")

##testing the fit of the models
library(AICcmodavg)

model1 <- lm(time ~ dat$`Mortality (%)` + formulation, data = dat)
model2 <- lm(time ~ dat$`Mortality (%)`, data = dat)

models <- list(model1, model2)
mod.names <- c('formulation', 'sans formulation')
```

```
aictab(cand.set = models, modnames = mod.names)

models <- list(nls_fit2, nls_fit3b, nls_fit4)

mod.names <- c('2 parameter', '3 parameter', '4 parameter')

aictab(cand.set = models, modnames = mod.names)
```
