## Supplemental Figure 1 for "Systematic review of residual toxicity studies of pesticides to bees and comparison to language on pesticide labels using data from studies and the Environmental Protection Agency"

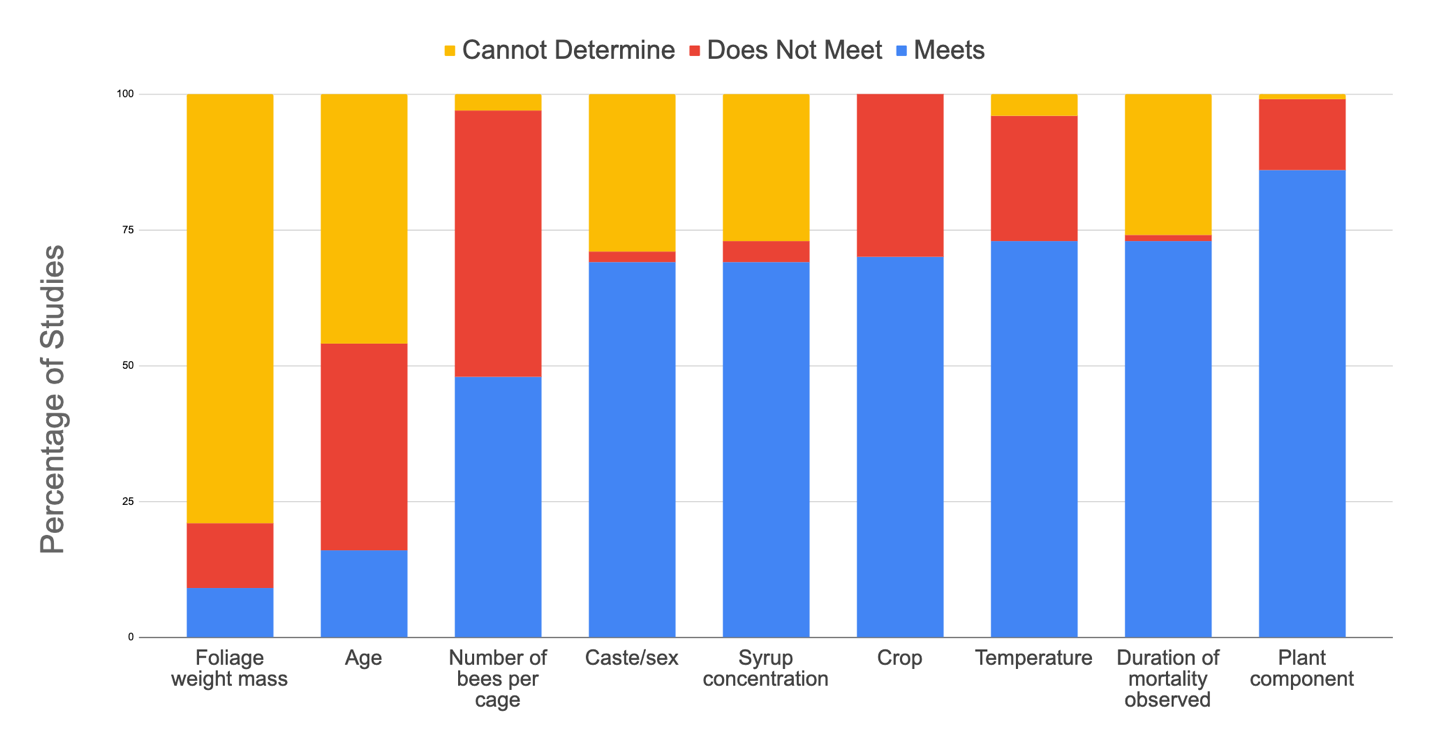

Figure 1: Comparison of methodological parameters of residual toxicity studies (n=48) with percentage of studies that meet USEPA residual toxicity criteria (USEPA, 2012a), do not meet, and cannot determine based on lack of information provided in the study.
