## Supplemental Figure 2 for "Systematic review of residual toxicity studies of pesticides to bees and comparison to language on pesticide labels using data from studies and the Environmental Protection Agency"

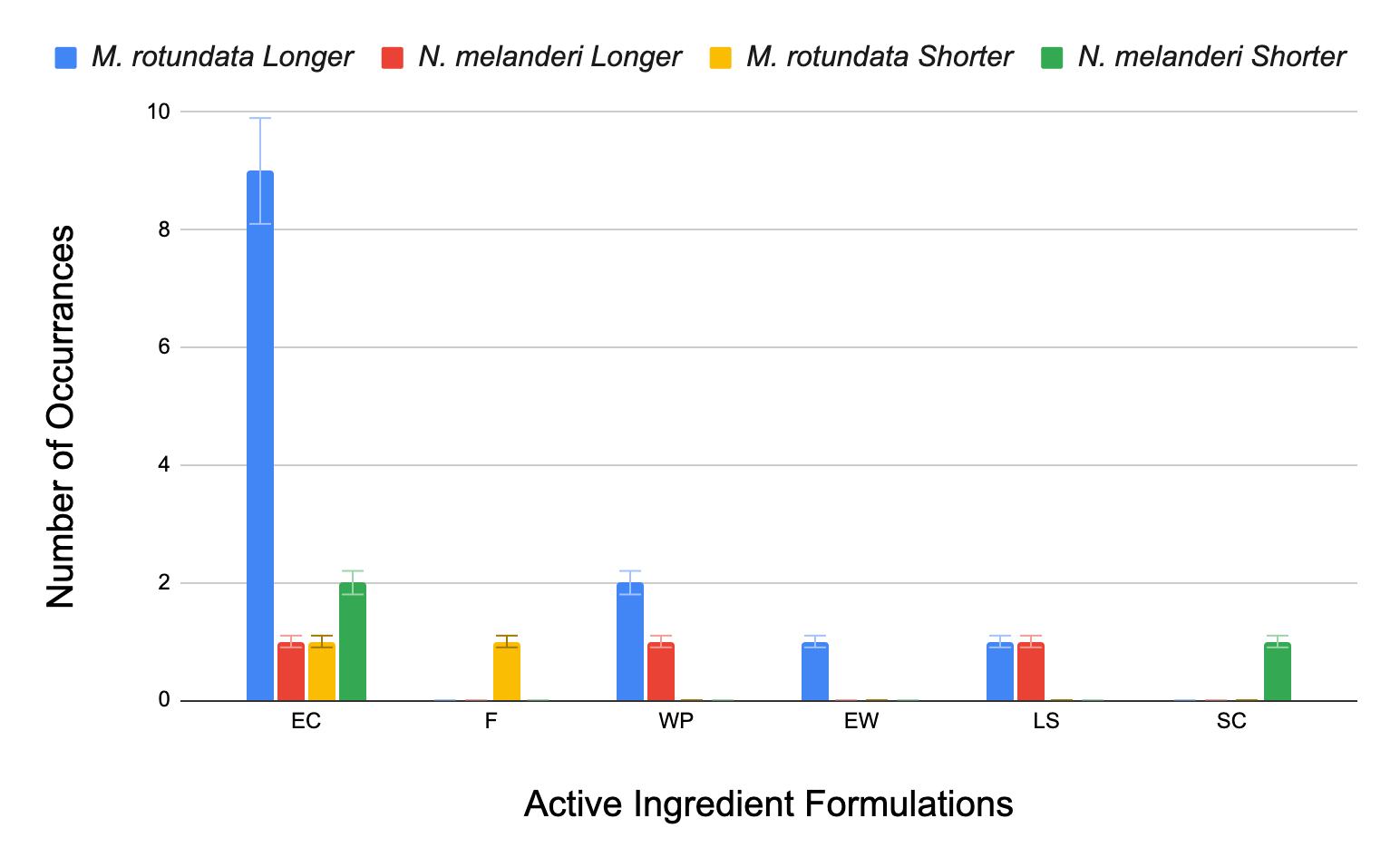

Figure 2: Comparison of bee species across different active ingredient formulations, EC (emulsifiable concentrate), F (flowable), WP (wettable powder), EW (emulsion in water), LS (liquid soluble), and SC (soluble concentrate. *M. rotundata* and *N. melanderi* RT_25_ values were compared to *A. mellifera* and reported as either (1) longer than *A. mellifera* values (“*M. rotundata* longer” and “*N. melanderi* longer”) or (2) shorter than *A. mellifera* values (“*M. rotundata* Shorter” and “*N. melanderi* Shorter”).
