## Supplemental Figure 3 for "Systematic review of residual toxicity studies of pesticides to bees and comparison to language on pesticide labels using data from studies and the Environmental Protection Agency"

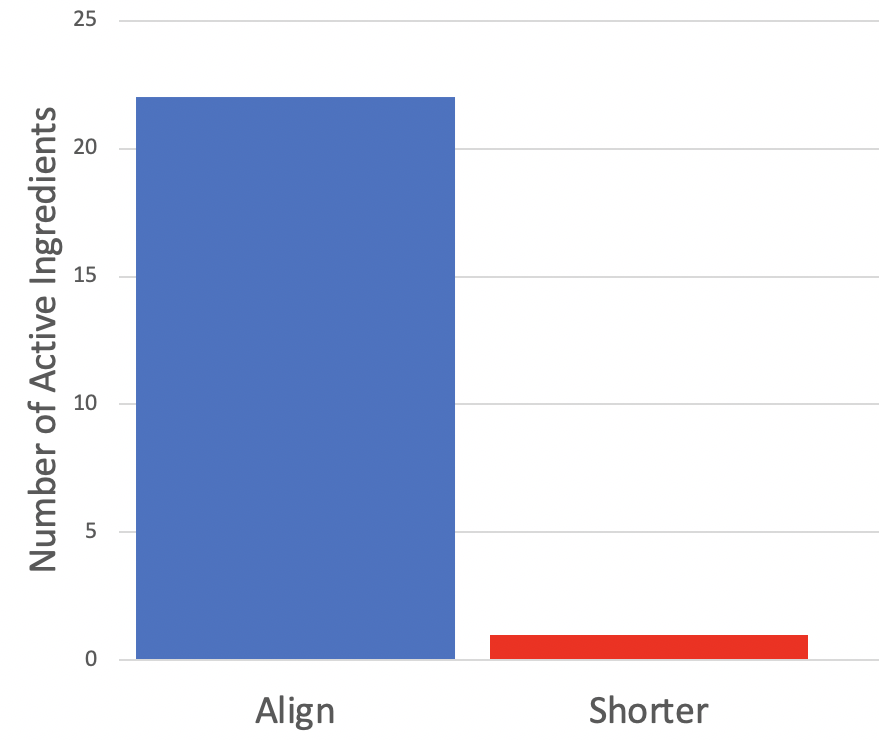

Figure 3: Total number of pesticide active ingredients where RT_25_ values calculated from the literature aligned in their approximations (*e.g*., both reported as greater than 24 hours) or reported the same value with the published by USEPA RT_25_ database (USEPA, 2014) signified by “Aligned”. If the literature and USEPA disagree on the RT_25_ it is depicted with the U reporting a “Shorter” RT_25_ value. We matched formulation and rate when comparing the USEPA values and the literature values.
