## Supplemental Figure 4 for "Systematic review of residual toxicity studies of pesticides to bees and comparison to language on pesticide labels using data from studies and the Environmental Protection Agency"

|  | Number of Active Ingredients |
| --- | --- |
| Literature | 29 |
| USEPA | 39 |
| Both | 30 |

Figure 4: Number of pesticide active ingredients where RT_25_ values could only be calculated from the literature (“Literature”), only from USEPA’s published database (“USEPA”; USEPA, 2014) or where there were RT_25_ values available from both sources (“Both”).
