## Supplemental Figure 5 for "Systematic review of residual toxicity studies of pesticides to bees and comparison to language on pesticide labels using data from studies and the Environmental Protection Agency"

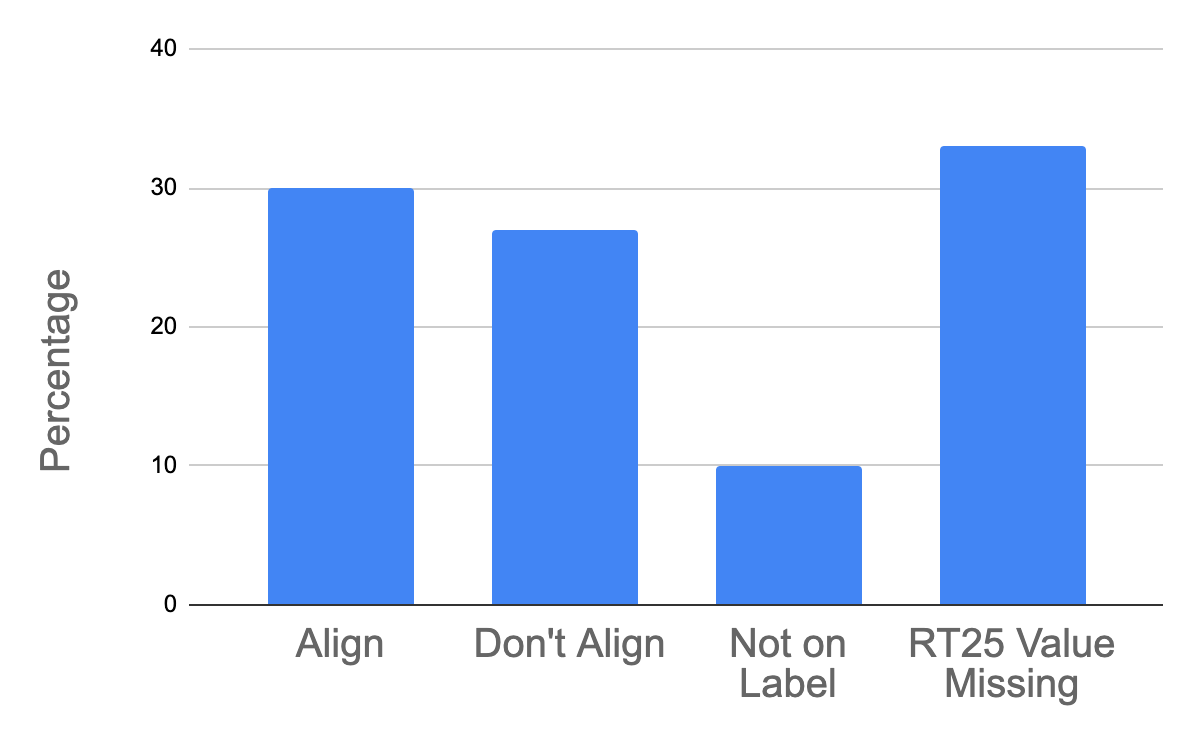

Figure 5: Comparison of pesticide label language indicating residual toxicity in relation to RT_25_ values (calculated from the literature and from USEPA (USEPA, 2014)). Residual toxicity language in the Environmental Hazards section either: (1) aligned with RT_25_ values (“Align”), (2) did not align (“Don’t Align”), (3) lacked residual toxicity language (“Not on Label”) or (4) did not have an RT_25_ value to relate to the label language (“RT_25_ value missing”). Formulation was matched when comparing label language to calculated RT_25_ values.
