## Supplemental Table 1 for "Systematic review of residual toxicity studies of pesticides to bees and comparison to language on pesticide labels using data from studies and the Environmental Protection Agency"

Table 1. Descriptions of Study Design Elements Examined during the Meta-Analysis

| **Study Design Element** | **Variables Examined** | **USEPA Guidelines** |
| --- | --- | --- |
| Bees^i^ | 1. Caste/sex 2. Age | 1. Female worker bees 2. Young |
| Plant Materials^ii^ | 1. Crop 2. Plant part | 1. Alfalfa 2. Foliage |
| Exposure^iii^ | 1. Number of bees per cage 2. Plant part weight mass 3. Duration of mortality observation | 1. 25 per cage 2. 15 grams 3. Greater than or equal to 24 hours |
| Environmental Conditions^iv^ | 1. Temperature 2. Syrup provided 3. Syrup concentration | 1. 25 to 35 degrees Celsius 2. Yes 3. 50:50 weight to volume |

^i^ Consists of the caste, sex, age, and source of bees placed in the cage during the residual toxicity trail. Caste = either worker, drone, or queen. Sex = either male or female. Age of bees = how old the bees (in days) generally were.

^ii^ Pertains to the materials used during the residual toxicity trials. Crop = the type of crop the product was sprayed on. Plant part = the part of the plan placed in the cages with the bees.

^iii^ How the bees were exposed to the pesticide. Number of bees per cage = the number of bees placed in each cage during the residual toxicity trial. Plant part weight mass = the weight mass of the plant part placed in the cage during the trial. Duration of mortality observation = how long in hours the bees were observed for mortality after being exposed.

^iv^ The environmental conditions that the bees were held at during the residual toxicity trial. Temperature = the average temperature the bees were incubated at during the trial. Syrup provided = if syrup was provided during the observation period. Syrup concentration = the concentration of the syrup in terms of water to sucrose.
