## Supplementary material for "Systematic review of residual toxicity studies of pesticides to bees and comparison to language on pesticide labels using data from studies and the Environmental Protection Agency": Supplmental Table 2

Table 2: Compiled Calculated RT_25_ database

| Active Ingredient | Formulation | Rate  (lb ai/A) | Greater than or less than 8 hours RT25 values | | | Test Species Scientific Name |
| --- | --- | --- | --- | --- | --- | --- |
|  |  |  | Meta-Analysis | # of studies | EPA Reported Value |  |
| Acephate | LS | 1 | > 24 | 1 |  | *Apis mellifera* |
|  | SP | 0.5 | > 24 | 1 |  | *Apis mellifera* |
|  |  |  | > 24 | 1 |  | *Megachile rotundata* |
|  |  | 1.29 | > 8 | 1 |  | *Apis mellifera* |
|  |  |  | > 8 | 1 |  | *Megachile rotundata* |
|  |  |  | > 8 | 1 |  | *Nomia melanderi* |
|  | WP | 1 | 7 or > 72 | 3 |  | *Apis mellifera* |
|  |  |  | 7.2 or > 72 | 3 |  | *Megachile rotundata* |
|  |  |  | 7.78 or > 72 | 3 |  | *Apis mellifera* |
| Acetamiprid | WP | 0.05 | < 2 | 1 |  | *Apis mellifera* |
|  |  |  | < 2 | 1 |  | *Megachile rotundata* |
|  |  |  | < 2 | 1 |  | *Nomia melanderi* |
|  |  | 0.075 | < 2 | 1 |  | *Apis mellifera* |
|  |  |  | < 2 | 1 |  | *Megachile rotundata* |
|  |  |  | < 2 | 1 |  | *Nomia melanderi* |
|  |  | 0.1 | < 2 | 1 |  | *Apis mellifera* |
|  |  |  | < 2 | 1 |  | *Megachile rotundata* |
|  |  |  | < 2 | 1 |  | *Nomia melanderi* |
|  |  | 0.15 | < 2 | 1 |  | *Apis mellifera* |
|  |  |  | < 2 | 1 |  | *Megachile rotundata* |
|  |  |  | > 8 | 1 |  | *Nomia melanderi* |
|  |  | 0.3 | < 2 | 1 |  | *Apis mellifera* |
|  |  |  | < 2 | 1 |  | *Megachile rotundata* |
|  |  |  | > 8 | 1 |  | *Nomia melanderi* |
| Aldoxycarb | F | 3 | > 8 | 1 |  | *Apis mellifera* |
| Azamethiphos | WP | 0.5 | > 144 | 1 |  | *Apis mellifera* |
|  |  | 2 | > 144 | 1 |  |  |
| Azinphos-methyl | EC | 1 | > 8 | 1 |  | *Apis mellifera* |
|  |  |  | > 8 | 1 |  | *Megachile rotundata* |
|  |  |  | > 8 | 1 |  | *Nomia melanderi* |
|  | WP | 1 | > 8 | 1 |  | *Apis mellifera* |
|  |  |  | > 8 | 1 |  | *Megachile rotundata* |
|  |  |  | > 8 | 1 |  | *Nomia melanderi* |
| Bifenthrin | E | 0.032 | 19 | 1 |  | *Apis mellifera* |
|  |  | 0.05 | > 24 | 1 |  | *Apis mellifera* |
|  | EC | 0.0125 | 63 | 1 |  | *Apis mellifera* |
|  |  |  | > 72 | 1 |  | *Megachile rotundata* |
|  |  | 0.025 | > 72 | 1 |  | *Apis mellifera* |
|  |  |  | > 72 | 1 |  | *Megachile rotundata* |
|  |  | 0.05 | > 72 | 1 |  | *Apis mellifera* |
|  |  |  | > 72 | 1 |  | *Megachile rotundata* |
|  |  | 0.06 | 128 | 1 |  | *Apis mellifera* |
|  |  | 0.1 | > 72 | 1 |  | *Apis mellifera* |
|  |  |  | > 72 | 1 |  | *Megachile rotundata* |
|  | ULV | 0.06 | 81.2 | 1 |  | *Apis mellifera* |
| Carbaryl | F | 3 | > 8 | 1 |  | *Apis mellifera* |
|  |  |  | > 8 | 1 |  | *Megachile rotundata* |
|  |  |  | > 8 | 1 |  | *Nomia melanderi* |
|  | WP | 0.25 | > 42 | 1 | > 42 | *Apis mellifera* |
|  |  | 0.5 | > 42 | 1 | > 42 |  |
|  |  | 1 | > 48 | 2 | > 42 | *Apis mellifera* |
|  |  |  | > 48 | 1 |  | *Megachile rotundata* |
|  |  |  | > 48 | 1 |  | *Nomia melanderi* |
|  |  | 2 | > 42 | 1 | > 42 | *Apis mellifera* |
| Carbofuran | F | 0.245 | > 8 | 1 |  | *Apis mellifera* |
|  |  |  | > 8 | 1 |  | *Megachile rotundata* |
|  |  |  | > 8 | 1 |  | *Nomia melanderi* |
|  |  | 1 | > 336 | 1 |  | *Apis mellifera* |
|  |  |  | 288 | 1 |  | *Megachile rotundata* |
|  |  |  | > 72 | 1 |  | *Nomia melanderi* |
| Chlorpyrifos | E | 0.75 | > 8 | 1 |  | *Apis mellifera* |
|  |  |  | > 8 | 1 |  | *Megachile rotundata* |
|  |  |  | > 8 | 1 |  | *Nomia melanderi* |
|  |  | 1 | > 12 | 1 |  | *Apis mellifera* |
|  |  | 1.5 | > 8 | 1 |  | *Apis mellifera* |
|  |  |  | > 8 | 1 |  | *Megachile rotundata* |
|  |  |  | > 8 | 1 |  | *Nomia melanderi* |
|  | EC | 0.025 | > 8 | 1 |  | *Apis mellifera* |
|  |  | 0.05 | > 8 | 1 |  | *Apis mellifera* |
|  |  | 0.1 | > 8 | 1 |  | *Apis mellifera* |
|  |  | 0.25 | 17 | 1 | 16 | *Apis mellifera* |
|  |  |  | > 24 | 1 | > 24 | *Megachile rotundata* |
|  |  |  | 20 | 1 | 19 | *Nomia melanderi* |
|  |  | 0.5 | 99 | 2 | > 24 | *Apis mellifera* |
|  |  |  | 140 | 2 | > 24 | *Megachile rotundata* |
|  |  |  | 66.8 | 2 | > 24 | *Nomia melanderi* |
|  |  | 1 | 141 | 2 | > 24 | *Apis mellifera* |
|  |  |  | 161 | 2 | > 24 | *Megachile rotundata* |
|  |  |  | > 120 | 2 | > 24 | *Nomia melanderi* |
| Clofentezine | F | 0.25 | < 2 | 1 |  | *Apis mellifera* |
|  |  |  | < 2 | 1 |  | *Megachile rotundata* |
|  |  | 0.5 | < 2 | 1 |  | *Apis mellifera* |
|  |  |  | < 2 | 1 |  | *Megachile rotundata* |
|  |  | 1 | < 2 | 1 |  | *Apis mellifera* |
|  |  |  | < 2 | 1 |  | *Megachile rotundata* |
|  |  | 2 | < 2 | 1 |  | *Apis mellifera* |
|  |  |  | < 2 | 1 |  | *Megachile rotundata* |
| Colpyralid | EC | 0.05 | < 2 | 1 |  | *Apis mellifera* |
|  |  |  | < 2 | 1 |  | *Megachile rotundata* |
|  |  | 0.1 | < 2 | 2 |  | *Apis mellifera* |
|  |  |  | < 2 | 1 |  | *Megachile rotundata* |
|  |  | 0.2 | 5 | 1 |  | *Megachile rotundata* |
|  | EW | 0.05 | < 2 | 1 |  | *Apis mellifera* |
|  |  |  | 3 | 1 |  | *Megachile rotundata* |
|  |  | 0.1 | < 2 | 3 |  | *Apis mellifera* |
|  |  |  | 44.6 | 2 |  | *Megachile rotundata* |
|  |  |  | < 8 | 1 |  | *Nomia melanderi* |
|  |  | 0.2 | < 2 | 1 |  | *Apis mellifera* |
|  |  |  | > 72 | 1 |  | *Megachile rotundata* |
| Cyfluthrin | E | 0.025 | > 24 | 1 |  | *Apis mellifera* |
|  |  |  | > 24 | 1 |  | *Megachile rotundata* |
|  |  | 0.05 | > 24 | 1 | > 240 | *Apis mellifera* |
|  |  |  | > 24 | 1 |  | *Megachile rotundata* |
| Cyhalothrin | E | 0.05 | > 8 | 1 |  | *Apis mellifera* |
|  |  |  | > 8 | 1 |  | *Megachile rotundata* |
|  |  |  | > 8 | 1 |  | *Nomia melanderi* |
|  | EC | 0.01 | < 2 | 1 |  | *Apis mellifera* |
|  |  |  | > 8 | 1 |  | *Megachile rotundata* |
|  |  |  | 2 | 1 |  | *Nomia melanderi* |
|  |  | 0.015 | < 2 | 1 |  | *Apis mellifera* |
|  |  |  | 8 | 1 |  | *Megachile rotundata* |
|  |  |  | 3.64 | 1 |  | *Nomia melanderi* |
|  |  | 0.02 | < 2 | 1 |  | *Apis mellifera* |
|  |  |  | > 8 | 1 |  | *Megachile rotundata* |
|  |  |  | 3.24 | 1 |  | *Nomia melanderi* |
|  |  | 0.025 | 4.28 | 1 |  | *Apis mellifera* |
|  |  |  | > 8 | 1 |  | *Megachile rotundata* |
|  |  |  | 6.54 | 1 |  | *Nomia melanderi* |
|  |  | 0.03 | > 8 | 1 |  | *Apis mellifera* |
|  |  |  | > 8 | 1 |  | *Megachile rotundata* |
|  |  |  | > 8 | 1 |  | *Nomia melanderi* |
| Cypermethrin | E | 0.05 | > 8 | 1 | > 96 | *Apis mellifera* |
|  |  |  | > 8 | 1 |  | *Megachile rotundata* |
|  |  |  | > 8 | 1 |  | *Nomia melanderi* |
|  |  | 0.1 | > 24 | 1 |  | *Apis mellifera* |
|  | EC | 0.06 | > 8 | 1 |  | *Apis mellifera* |
|  |  | 0.09 | 313 | 1 |  | *Megachile rotundata* |
|  |  | 0.14 | 197 | 1 |  | *Apis mellifera* |
|  | ULV | 0.09 | 63.8 | 1 |  | *Apis mellifera* |
| Cyromazine | WP | 025 | < 2 | 1 |  | *Apis mellifera* |
|  |  | 0.3 | > 8 | 1 |  | *Megachile rotundata* |
|  |  |  | > 8 | 1 |  | *Nomia melanderi* |
| Deltamethrin | EC | 0.02 | 4.95 | 1 | 5.2 | *Apis mellifera* |
|  |  | 0.2 | < 2 | 1 |  | *Apis mellifera* |
|  |  |  | 4.09 | 1 |  | *Megachile rotundata* |
|  |  |  | < 2 | 1 |  | *Nomia melanderi* |
| Diazinon | EC | 0.05 | > 24 | 1 |  | *Apis mellifera* |
|  |  |  | > 8 | 1 |  | *Megachile rotundata* |
|  |  | 0.75 | > 8 | 1 |  | *Apis mellifera* |
|  |  |  | > 8 | 1 |  | *Megachile rotundata* |
|  |  |  | > 8 | 1 |  | *Nomia melanderi* |
|  |  | 1.5 | > 8 | 1 |  | *Apis mellifera* |
|  |  |  | > 8 | 1 |  | *Megachile rotundata* |
|  |  |  | > 8 | 1 |  | *Nomia melanderi* |
|  |  | 3 | > 8 | 1 |  | *Apis mellifera* |
|  |  |  | > 8 | 1 |  | *Megachile rotundata* |
|  |  |  | > 8 | 1 |  | *Nomia melanderi* |
|  | WP | 0.125 | > 18 | 1 | < 42 | *Apis mellifera* |
|  |  | 0.25 | > 18 | 1 | < 42 |  |
|  |  | 0.5 | > 42 | 1 | > 42 |  |
|  |  | 1 | > 42 | 1 | > 42 |  |
| Dicofol | EC | 1.5 | < 3 | 1 |  | *Apis mellifera* |
| Dimethoate | EC | 0.125 | < 3 | 1 |  | *Apis mellifera* |
|  |  |  | < 3 | 1 |  | *Megachile rotundata* |
|  |  | 0.25 | 4.18 | 1 |  | *Apis mellifera* |
|  |  |  | 3 | 1 |  | *Megachile rotundata* |
|  |  | 0.5 | 114 or 11.9 | 2 | < 120 | *Apis mellifera* |
|  |  |  | 121 | 2 | < 120 | *Megachile rotundata* |
|  |  |  | > 72 | 2 | > 72 | *Nomia melanderi* |
| Disulfoton | EC | 0.5 | < 3 | 1 |  | *Apis mellifera* |
|  |  |  | 13 | 1 |  | *Megachile rotundata* |
|  |  |  | < 3 | 1 |  | *Nomia melanderi* |
|  |  | 1 | 8.86 | 1 | 5.5 | *Apis mellifera* |
|  |  |  | 20.7 | 1 |  | *Megachile rotundata* |
|  |  |  | 2.23 | 1 |  | *Nomia melanderi* |
| Endosulfan | EC | 0.75 | < 2 | 1 | < 3 | *Apis mellifera* |
|  |  |  | > 8 | 1 |  | *Megachile rotundata* |
|  |  |  | 6.75 | 1 |  | *Nomia melanderi* |
|  | WP | 0.5 | < 8 | 1 |  | *Apis mellifera* |
|  |  |  | > 8 | 1 |  | *Megachile rotundata* |
|  |  | 0.75 | > 8 | 1 |  | *Megachile rotundata* |
|  |  | 1 | > 8 | 1 |  |  |
| Esfenvalerate | EC | 0.0125 | < 2 | 1 |  | *Apis mellifera* |
|  |  | 0.05 | > 8 | 2 |  | *Apis mellifera* |
|  |  |  | < 2 | 2 |  | *Megachile rotundata* |
|  |  |  | 8 | 1 |  | *Nomia melanderi* |
|  |  | 0.075 | > 8 | 1 |  | *Apis mellifera* |
|  |  |  | < 2 | 1 |  | *Megachile rotundata* |
|  |  | 0.1 | > 24 | 2 |  | *Apis mellifera* |
|  |  |  | < 2 | 1 |  | *Megachile rotundata* |
| Ethiprole | EC | 0.18 | 643 | 1 |  | *Apis mellifera* |
|  | SC | 0.3 | 333 | 1 |  |  |
| Fenitrothion | EC | 0.5 | 18.2 | 2 | < 24 | *Apis mellifera* |
|  |  |  | > 72 | 1 | 106 | *Megachile rotundata* |
|  |  |  | > 72 | 1 | 98 | *Nomia melanderi* |
|  |  | 1 | > 72 | 2 | 101 | *Apis mellifera* |
|  |  |  | > 120 | 1 | > 120 | *Megachile rotundata* |
|  |  |  | > 120 | 1 | > 120 | *Nomia melanderi* |
| Fenpropathrin | EC | 0.1 | > 8 | 1 | < 192 | *Apis mellifera* |
|  |  |  | > 8 | 1 |  | *Megachile rotundata* |
|  |  |  | > 8 | 1 |  | *Nomia melanderi* |
|  |  | 0.2 | > 8 | 2 | 276 | *Apis mellifera* |
|  |  |  | > 8 | 1 |  | *Megachile rotundata* |
|  |  |  | > 8 | 1 |  | *Nomia melanderi* |
|  |  | 0.4 | > 8 | 2 | < 336 | *Apis mellifera* |
|  |  |  | > 8 | 1 |  | *Megachile rotundata* |
|  |  |  | > 8 | 1 |  | *Nomia melanderi* |
| Fenvalerate | EC | 0.1 | 6.5 | 2 | 7 | *Apis mellifera* |
|  |  |  | > 8 | 1 | > 8 | *Megachile rotundata* |
|  |  |  | 6.82 | 2 | 7 | *Nomia melanderi* |
|  |  | 0.2 | 16.4 | 1 |  | *Apis mellifera* |
|  |  |  | > 8 | 1 |  | *Megachile rotundata* |
|  |  | 0.4 | > 8 | 1 | > 8 | *Apis mellifera* |
|  |  |  | > 8 | 1 | > 8 | *Megachile rotundata* |
|  |  |  | > 8 | 1 | > 8 | *Nomia melanderi* |
| Fipronil | SC | 0.01 | 238 | 1 |  | *Apis mellifera* |
|  |  | 0.0125 | < 2 | 1 |  | *Apis mellifera* |
|  |  |  | 3.82 | 1 |  | *Megachile rotundata* |
|  |  |  | < 2 | 1 |  | *Nomia melanderi* |
|  |  | 0.025 | < 2 | 1 |  | *Apis mellifera* |
|  |  |  | < 2 | 1 |  | *Megachile rotundata* |
|  |  |  | < 2 | 1 |  | *Nomia melanderi* |
|  |  | 0.1 | 7.15 | 1 |  | *Apis mellifera* |
|  |  |  | > 8 | 1 |  | *Megachile rotundata* |
|  |  |  | < 2 | 1 |  | *Nomia melanderi* |
|  |  | 0.2 | > 8 | 1 |  | *Apis mellifera* |
|  |  |  | > 8 | 1 |  | *Megachile rotundata* |
|  |  |  | < 2 | 1 |  | *Nomia melanderi* |
|  | WG | 0.0125 | < 2 | 1 |  | *Apis mellifera* |
|  |  |  | < 2 | 1 |  | *Megachile rotundata* |
|  |  |  | < 2 | 1 |  | *Nomia melanderi* |
|  |  | 0.025 | < 2 | 1 |  | *Apis mellifera* |
|  |  |  | < 2 | 1 |  | *Megachile rotundata* |
|  |  |  | < 2 | 1 |  | *Nomia melanderi* |
|  |  | 0.1 | 5.51 or > 8 | 2 |  | *Apis mellifera* |
|  |  |  | > 8 or 3.52 | 2 |  | *Megachile rotundata* |
|  |  |  | < 2 | 2 |  | *Nomia melanderi* |
|  |  | 0.2 | > 8 | 2 |  | *Apis mellifera* |
|  |  |  | > 8 | 1 |  | *Megachile rotundata* |
|  |  |  | < 2 | 1 |  | *Nomia melanderi* |
| Fluazinam | WDG | 0.135 | < 2 | 1 |  | *Megachile rotundata* |
| Flupyradifurone | SL | 0.183 | < 3 | 1 | < 3 | *Apis mellifera* |
| Fluvalinate | E | 0.1 | < 2 | 1 |  | *Apis mellifera* |
| Fonofos | Enc. | 1 | < 3 | 1 |  | *Apis mellifera* |
|  |  | 2 | > 8 | 1 |  |  |
|  | EC | 1 | < 3 | 1 | < 3 |  |
|  |  | 2 | 5.76 | 1 | < 8 |  |
| Formetanate Hydrochloride | SP | 0.23 | < 3 | 1 |  | *Apis mellifera* |
|  |  |  | < 3 | 1 |  | *Megachile rotundata* |
|  |  |  | < 3 | 1 |  | *Nomia melanderi* |
|  |  | 0.45 | < 3 | 1 |  | *Apis mellifera* |
|  |  |  | < 3 | 1 |  | *Megachile rotundata* |
|  |  |  | < 3 | 1 |  | *Nomia melanderi* |
|  |  | 0.5 | < 2 | 4 |  | *Apis mellifera* |
|  |  |  | < 3, 7.5, or > 8 | 4 |  | *Megachile rotundata* |
|  |  |  | 11.2 or < 3 | 3 |  | *Nomia melanderi* |
|  |  | 1 | 4.32 | 3 |  | *Apis mellifera* |
|  |  |  | 5.3 | 2 |  | *Megachile rotundata* |
|  |  |  | 5.15 | 1 |  | *Nomia melanderi* |
|  |  | 1.1 | 6.68 | 1 |  | *Apis mellifera* |
|  |  |  | > 8 | 1 |  | *Megachile rotundata* |
|  |  |  | > 8 | 1 |  | *Nomia melanderi* |
| Imidacloprid | EC | 0.25 | 90 | 1 |  | *Apis mellifera* |
|  |  |  | 214 | 1 |  | *Megachile rotundata* |
|  |  |  | > 72 | 1 |  | *Nomia melanderi* |
|  |  | 0.5 | 110 | 1 |  | *Apis mellifera* |
|  |  |  | 277 | 2 |  | *Megachile rotundata* |
|  |  |  | > 72 | 1 |  | *Nomia melanderi* |
|  | F | 0.15 | < 2 | 1 |  | *Apis mellifera* |
|  |  |  | > 8 | 1 |  | *Megachile rotundata* |
|  |  |  | 2.72 | 1 |  | *Nomia melanderi* |
|  |  | 0.1 | 2.56 | 1 | < 8 | *Nomia melanderi* |
|  | SL | 0.018 | 236 | 1 |  | *Apis mellifera* |
|  | WG | 0.045 | < 3 | 1 |  | *Apis mellifera* |
|  |  | 0.167 | 31.1 | 1 |  |  |
|  |  | 0.5 | 89.8 | 1 |  |  |
| Indoxacarb | SC | 0.039 | 140 | 1 |  | *Apis mellifera* |
| Lambda-cyhalothrin | E | 0.02 | 17 | 1 |  | *Apis mellifera* |
|  |  | 0.03 | > 24 | 1 |  | *Apis mellifera* |
|  |  |  | > 8 | 1 |  | *Megachile rotundata* |
|  | EC | 0.01 | 54 | 1 |  | *Apis mellifera* |
|  |  |  | > 72 | 1 |  | *Megachile rotundata* |
|  |  | 0.02 | > 72 | 1 |  | *Apis mellifera* |
|  |  |  | > 72 | 1 |  | *Megachile rotundata* |
| Leptophos | EC | 1 | 2.32 | 2 |  | *Apis mellifera* |
|  |  |  | 13.8 | 2 |  | *Megachile rotundata* |
|  |  |  | 3.86 | 2 |  | *Nomia melanderi* |
|  |  | 2 | > 8 | 1 |  | *Apis mellifera* |
| Lindane | EC | 0.5 | > 8 | 1 | 24 | *Apis mellifera* |
|  |  | 1 | > 24 | 1 | 72 |  |
|  |  | 1.5 | > 48 | 1 | 72 |  |
|  | F | 0.5 | > 8 | 1 | 24 |  |
|  |  | 1 | > 48 | 1 | 72 |  |
|  |  | 1.5 | > 72 | 1 | 72 |  |
|  | WP | 0.5 | > 8 | 1 | 24 |  |
|  |  | 1 | > 48 | 1 | 72 |  |
|  |  | 1.5 | > 48 | 1 | 72 |  |
| Malathion | E | 1 | > 8 | 1 |  | *Apis mellifera* |
|  |  |  | > 8 | 1 |  | *Megachile rotundata* |
|  |  |  | > 8 | 1 |  | *Nomia melanderi* |
|  | EC | 0.625 | > 18 | 1 |  | *Apis mellifera* |
|  |  | 1 | > 24 | 1 |  |  |
|  |  | 1.25 | > 42 | 1 |  |  |
|  | WP | 0.3125 | > 18 | 1 |  | *Apis mellifera* |
|  |  | 0.625 | > 42 | 1 |  |  |
|  |  | 1.25 | > 42 | 1 |  |  |
| Malonoben | EC | 0.5 | < 8 | 1 |  | *Megachile rotundata* |
|  |  |  | < 2 | 1 |  | *Nomia melanderi* |
|  |  | 1 | > 8 | 1 |  | *Megachile rotundata* |
|  |  |  | < 2 | 1 |  | *Nomia melanderi* |
|  |  | 2 | > 24 | 1 |  | *Megachile rotundata* |
|  |  |  | > 8 | 1 |  | *Nomia melanderi* |
|  | WP | 0.25 | < 2 | 1 |  | *Apis mellifera* |
|  |  |  | < 2 | 1 |  | *Megachile rotundata* |
|  |  |  | < 2 | 1 |  | *Nomia melanderi* |
|  |  | 0.5 | < 2 | 1 |  | *Apis mellifera* |
|  |  |  | 6 | 1 |  | *Megachile rotundata* |
|  |  |  | < 2 | 1 |  | *Nomia melanderi* |
|  |  | 1 | < 2 | 1 |  | *Apis mellifera* |
|  |  |  | 18 | 1 |  | *Megachile rotundata* |
|  |  |  | < 2 | 1 |  | *Nomia melanderi* |
| Methamidophos | EC | 0.67 | > 8 | 1 |  | *Apis mellifera* |
|  |  |  | > 8 | 1 |  | *Megachile rotundata* |
|  |  |  | > 8 | 1 |  | *Nomia melanderi* |
| Methidathion | E | 0.736 | > 8 | 1 |  | *Apis mellifera* |
|  |  |  | > 8 | 1 |  | *Megachile rotundata* |
|  |  |  | > 8 | 1 |  | *Nomia melanderi* |
|  | EC | 1 | 91 | 1 |  | *Apis mellifera* |
|  |  |  | 89.6 | 1 |  | *Megachile rotundata* |
|  |  |  | > 72 | 1 |  | *Nomia melanderi* |
| Methomyl | EC | 0.9 | < 2 | 1 |  | *Apis mellifera* |
|  | LS | 0.25 | < 3 | 1 |  | *Apis mellifera* |
|  |  |  | < 4 | 1 |  | *Megachile rotundata* |
|  |  |  | < 4 | 1 |  | *Nomia melanderi* |
|  |  | 0.5 | < 3 | 1 |  | *Apis mellifera* |
|  |  |  | < 4 | 1 |  | *Megachile rotundata* |
|  |  |  | 5 | 1 |  | *Nomia melanderi* |
|  |  | 1 | 6.11 | 1 |  | *Apis mellifera* |
|  |  |  | 20.5 | 1 |  | *Megachile rotundata* |
|  |  |  | > 24 | 1 |  | *Nomia melanderi* |
|  | WP | 0.5 | < 2 | 1 |  | *Apis mellifera* |
|  |  |  | 5.2 | 1 |  | *Megachile rotundata* |
|  |  |  | 4.53 | 1 |  | *Nomia melanderi* |
|  |  | 0.9 | > 8 | 1 |  | *Apis mellifera* |
|  |  |  | > 8 | 1 |  | *Megachile rotundata* |
|  |  |  | > 8 | 1 |  | *Nomia melanderi* |
|  |  | 1 | < 8 | 1 |  | *Apis mellifera* |
|  |  |  | 5.87 | 1 |  | *Megachile rotundata* |
|  |  |  | 6 | 1 |  | *Nomia melanderi* |
| Methyl Parathion | CS | 0.401 | 205 | 1 | 207 | *Apis mellifera* |
|  | EC | 0.5 | 76 | 3 |  | *Apis mellifera* |
|  |  |  | > 72 | 2 |  | *Megachile rotundata* |
|  |  |  | > 8 | 1 |  | *Nomia melanderi* |
|  |  | 1 | 81 | 2 |  | *Apis mellifera* |
|  |  |  | > 72 | 1 |  | *Megachile rotundata* |
|  | F | 0.5 | > 8 | 1 |  | *Apis mellifera* |
|  |  |  | > 8 | 1 |  | *Megachile rotundata* |
|  |  |  | > 8 | 1 |  | *Nomia melanderi* |
| Naled | E | 1 | > 8 or < 8 | 2 |  | *Apis mellifera* |
|  |  |  | > 8 | 1 |  | *Megachile rotundata* |
|  |  |  | 2 | 1 |  | *Nomia melanderi* |
|  | EC | 1 | > 8 | 2 |  | *Apis mellifera* |
|  |  |  | 6.44 or > 72 | 2 |  | *Megachile rotundata* |
|  |  |  | > 24 | 1 |  | *Nomia melanderi* |
| Oxamyl | EC | 1 | > 24 | 1 |  | *Apis mellifera* |
|  | LS | 0.25 | < 4 | 1 |  | *Apis mellifera* |
|  |  |  | < 4 | 1 |  | *Megachile rotundata* |
|  |  |  | < 4 | 1 |  | *Nomia melanderi* |
|  |  | 0.5 | < 4 | 1 |  | *Apis mellifera* |
|  |  |  | > 9 | 1 |  | *Megachile rotundata* |
|  |  |  | > 9 | 1 |  | *Nomia melanderi* |
|  |  | 1 | 12.5 | 1 | 22 | *Apis mellifera* |
|  |  |  | > 24 | 1 |  | *Megachile rotundata* |
|  |  |  | > 24 | 1 |  | *Nomia melanderi* |
| Oxydemeton-methyl | EC | 0.5 | < 2 | 1 |  | *Apis mellifera* |
|  |  |  | < 2 | 1 |  | *Megachile rotundata* |
|  |  | 0.75 | < 2 | 1 |  | *Apis mellifera* |
|  |  |  | < 2 | 1 |  | *Megachile rotundata* |
|  |  | 1 | 6 | 1 |  | *Nomial melanderi* |
|  | SC | 0.5 | < 2 | 1 |  | *Apis mellifera* |
|  |  |  | < 2 | 1 |  | *Megachile rotundata* |
|  |  |  | < 2 | 1 |  | *Nomia melanderi* |
| Parathion | EC | 0.5 | 12.6 | 1 |  | *Apis mellifera* |
|  |  |  | 11.5 | 1 |  | *Megachile rotundata* |
|  |  |  | 12.8 | 1 |  | *Nomia melanderi* |
| Permethrin | EC | 0.05 | 21 | 1 |  | *Apis mellifera* |
|  |  |  | > 24 | 1 |  | *Megachile rotundata* |
|  |  |  | 15 | 1 |  | *Nomia melanderi* |
|  |  | 0.1 | 169 | 3 |  | *Apis mellifera* |
|  |  |  | > 24 | 1 |  | *Megachile rotundata* |
|  |  |  | > 24 | 1 |  | *Nomia melanderi* |
|  |  | 0.125 | > 8 | 1 |  | *Megachile rotundata* |
|  |  | 0.2 | > 168 | 2 |  | *Apis mellifera* |
|  |  |  | > 24 | 1 |  | *Megachile rotundata* |
|  |  |  | > 24 | 1 |  | *Nomia melanderi* |
|  | ULV | 0.1 | 95.3 | 1 |  | *Apis mellifera* |
|  | WP | 0.05 | > 72 | 1 |  | *Apis mellifera* |
|  |  |  | > 72 | 1 |  | *Megachile rotundata* |
|  |  | 0.1 | > 72 | 1 |  | *Apis mellifera* |
|  |  |  | > 72 | 1 |  | *Megachile rotundata* |
| Phenthoate | EC | 0.15625 | 18 | 1 |  | *Apis mellifera* |
|  |  | 0.3125 | > 18 | 1 |  |  |
|  |  | 0.625 | > 42 | 1 |  |  |
|  |  | 1.25 | > 42 | 1 |  |  |
| Phosmet | EC | 1 | > 8 | 1 |  | *Apis mellifera* |
|  |  | 2 | > 8 | 1 |  |  |
|  | WP | 1 | > 8 | 1 | > 3 |  |
|  |  | 2 | > 8 | 1 |  |  |
| Prochloraz | EC | 0.5 | < 2 | 1 |  | *Apis mellifera* |
|  |  | 1 | < 2 | 1 |  |  |
|  |  | 2 | < 2 | 1 |  |  |
| Profenofos | EC | 1 | > 8 | 1 |  | *Apis mellifera* |
|  |  |  | > 8 | 1 |  | *Megachile rotundata* |
|  |  |  | > 8 | 1 |  | *Nomia melanderi* |
| Propargite | EC | 2.1 | < 3 | 1 |  | *Apis mellifera* |
|  |  |  | < 3 | 1 |  | *Megachile rotundata* |
|  |  | 2.25 | < 3 | 1 |  | *Apis mellifera* |
|  |  |  | < 3 | 1 |  | *Megachile rotundata* |
| Piperonyl butoxide | E | 0.5 | > 24 | 1 |  | *Apis mellifera* |
| Pyrethrins | EC | 1 | < 2 | 1 |  | *Apis mellifera* |
|  |  |  | < 2 | 1 |  | *Megachile rotundata* |
|  |  |  | < 2 | 1 |  | *Nomia melanderi* |
| Sulfloxaflor | SC | 0.18 | < 1 | 3 |  | *Megachile rotundata* |
|  |  |  | < 1 | 3 |  | *Nomia melanderi* |
| Tetraniliprole | SC | 0.027 | < 3 | 1 |  | *Apis mellifera* |
|  |  | 0.054 | < 3 | 1 |  |  |
|  |  | 0.089 | < 3 | 1 |  |  |
| Thiacloprid | SC | 0.045 | < 2 | 1 |  | *Apis mellifera* |
|  |  | 0.09 | < 2 | 1 |  |  |
|  |  | 0.16 | < 2 | 1 | < 2 |  |
| Thiodicarb | F | 0.5 | < 2 | 1 |  | *Apis mellifera* |
|  |  | 1.2 | 77 | 1 |  |  |
|  | WDG | 1 | > 8 | 1 |  | *Apis mellifera* |
| Tiazamate | E | 0.25 | < 2 | 1 |  | *Apis mellifera* |
|  |  |  | < 2 | 1 |  | *Nomia melanderi* |
| Tolfenpyrad | EC | 1.69 | > 168 | 1 |  | *Megachile rotundata* |
|  |  |  | > 168 | 1 |  | *Nomia melanderi* |
| Trichlorfon | SP | 1 | < 8, or > 8 or 5.39 | 5 |  | *Apis mellifera* |
|  |  |  | 4.45 | 3 |  | *Megachile rotundata* |
|  |  |  | 4.64 | 2 |  | *Nomia melanderi* |
| Zeta-cypermethrin | EW | 0.037 | > 8 | 1 |  | *Apis mellifera* |
|  |  |  | > 8 | 1 |  | *Megachile rotundata* |
|  |  |  | > 8 | 1 |  | *Nomia melanderi* |
|  | WP | 1 | > 72 | 1 |  | *Apis mellifera* |
|  |  |  | > 8 | 1 |  | *Megachile rotundata* |
|  |  |  | > 8 | 1 |  | *Nomia melanderi* |

Key: E: emulsifiable, EC: emulsifiable concentrate, Enc.: encapsulated, EW: emulsion in water, F: flowable, LS: liquid soluble, SC: soluble concentrate, SL: soluble (liquid) concentrate, SP: soluble powder, ULV: ultra-low volume liquid, WDG: water dispersible granular, WG: wettable granule, WP: wettable powder
