## Supplemental Table 3 for "Systematic review of residual toxicity studies of pesticides to bees and comparison to language on pesticide labels using data from studies and the Environmental Protection Agency"

Table 3: Mode of Action Based on Availability in Insect Resistance Action Committee (2022) for all Active Ingredients in RT_25_ Database (Calculated and USEPA (2014)).

| Mode Action | Active Ingredients |
| --- | --- |
| 1A | Aldoxycarb, Bendiocarb, Carbaryl, Carbofuran, Formetanate Hydrochloride, Methomyl, Oxamyl, Propoxur, Thiodicarb, Tiazamate. |
| 1B | Acephate, Azemethiphos, Azinphos-methyl, Chlorpyrifos, Diazinon, Dichlorvos, Dicrotophos, Dimethoate, Disulfoton, Fenitrothion, Malathion, Methamidophos, Methidathion, Methyl parathion, Mevinphos, Naled, Oxydemeton-methyl, Parathion, Phenthoate, Phosalone, Phosmet, Profenofos, Trichlorfon |
| 2A | Endosulfan |
| 2B | Ethiprole, Fipronil |
| 3A | Bifenthrin, Cyfluthrin, Cyhalothrin, Cypermethrin, Deltamethrin, D-phenothrin, Esfenvalerate, Etofenprox, Fenpropathrin, Fenvalerate, Fluvalinate, Gamma cyhalothrin, Imiprothrin, Lambda-cyhalothrin, Momfluorothrin, Permethrin, Prallethrin, Pyrethrins, Resmethrin, Tetramethrin, Tralomethrin, Zeta-cypermethrin |
| 4A | Acetamiprid, Clothianidin, Dinotefuran, Imidacloprid, Thiacloprid, Thiamethoxam |
| 4C | Sulfloxaflor |
| 4D | Flupyradifuron |
| 5 | Spinetoram, Spinosad |
| 6 | Abamectin, Avermectin, Emamectin benozoate |
| 9B | Pyrifluquinazon |
| 9D | Afidopyropen |
| 10A | Clofentezine |
| 12B | Fenbutatin oxide |
| 12C | Propargite |
| 13 | Chlorfenapyr |
| 17 | Cyromazine |
| 19 | Amitraz |
| 20D | Bifenazate |
| 21A | Fenazaquin, Pyridaben, Tolfenpyrad |
| 22A | Indoxacarb |
| 28 | Chlorantraniliprole, Cyantraniliprole, Cyclaniliprole |
| UN | Azadirachtin, Dicofol |
