## Supplementary material for "Systematic review of residual toxicity studies of pesticides to bees and comparison to language on pesticide labels using data from studies and the Environmental Protection Agency": PRISMA Flow Diagram

**Identification of new studies via other methods**

**Previous studies**

**Identification of new studies via databases and registers**

Studies included in previous version of review (n = 0)

Reports of studies included in previous version of review (n = 0)

Records identified from*:

Databases (n = 130)

Registers (n = 0)

Records removed *before screening*:

Duplicate records removed

(n = 0)

Records marked as ineligible by automation tools (n = 0)

Records removed for other reasons (n = 0)

Records identified from:

Websites (n = 0)

Organisations (n = 53)

Citation searching (n = 0)

etc.

**Identification**

Reports not retrieved

(n = 0)

Reports sought for retrieval

(n = 0)

Total studies included in review

(n = )

Reports of total included studies

(n = )

Reports assessed for eligibility

(n = 4)

Reports sought for retrieval

(n = 0)

Records screened

(n = 130)

Records excluded**

(n = 126)

Reports excluded:

Reason 1: The study was not a primary source of data.

Reason 2: Bees were exposed to the pesticide applied to paper instead of on plant material.

Reason 3: The study focused on residual toxicity of pesticides applied to plots but did not involve harvesting plant tissue for caged bees.

Reports assessed for eligibility

(n = 53)

Reports not retrieved

(n = 0)

**Screening**

Reports excluded:

Reason 1: The study was not a primary source of data.

Reason 2: Bees were exposed to the pesticide applied to paper instead of on plant material.

Reason 3: The study focused on residual toxicity of pesticides applied to plots but did not involve harvesting plant tissue for caged bees.

New studies included in review

(n = 50)

Reports of new included studies

(n = )

**Included**

*Consider, if feasible to do so, reporting the number of records identified from each database or register searched (rather than the total number across all databases/registers).

**If automation tools were used, indicate how many records were excluded by a human and how many were excluded by automation tools.

From: Page MJ, McKenzie JE, Bossuyt PM, Boutron I, Hoffmann TC, Mulrow CD, et al. The PRISMA 2020 statement: an updated guideline for reporting systematic reviews. BMJ 2021;372:n71. doi: 10.1136/bmj.n71. For more information, visit: <http://www.prisma-statement.org/>
